# Epicardial therapy with atrial appendage micrografts salvages myocardium after infarction

**DOI:** 10.1101/712778

**Authors:** Xie Yanbo, Milla Lampinen, Juuso Takala, Vilbert Sikorski, Rabah Soliymani, Miikka Tarkia, Maciej Lalowski, Eero Mervaala, Markku Kupari, Zhe Zheng, Shengshou Hu, Ari Harjula, Esko Kankuri, on behalf of the AADC consortium

## Abstract

Ischemic heart disease remains the leading cause of mortality and morbidity worldwide despite improved possibilities in medical care. Alongside interventional therapies, such as coronary artery bypass grafting, adjuvant tissue-engineered and cell-based treatments can provide regenerative improvement. Unfortunately, most of these advanced approaches require multiple lengthy and costly preparation stages without delivering significant clinical benefits.

We evaluated the effect of epicardially delivered minute pieces of atrial appendage tissue material, defined as atrial appendage micrografts (AAMs), in mouse myocardial infarction model. An extracellular matrix patch was used to cover and fix the AAMs onto the surface of the infarcted heart. The matrix-covered AAMs salvaged the heart from infarction-induced loss of functional myocardium and attenuated scarring. Site-selective proteomics of injured ischemic and uninjured distal myocardium from AAM-treated and untreated tissue sections revealed an increased expression of several cardiac regeneration-associated proteins (i.e. periostin, transglutaminases and glutathione peroxidases) as well as activation of pathways responsible for angio- and cardiogenesis in relation to AAMs therapy.

Epicardial delivery of AAMs encased in an extracellular matrix patch scaffold salvages functional cardiac tissue from ischemic injury and restricts fibrosis after myocardial infarction. Our results support the use of AAMs as tissue-based therapy adjuvants for salvaging the ischemic myocardium.

## Introduction

Despite preventive efforts and increasing awareness of risk factors, ischemic heart disease persistently maintains its rank as the global leading cause of mortality (1). Irreversible tissue damage and death of functional myocardium after, for example a myocardial infarction (MI), is repaired by fibrotic scar contributing to loss-of-function, tissue rigidity and development of heart failure (HF) over time (2). Although drug therapies alleviate symptoms, they fail to restore lost myocardial tissue (3, 4). Hope has been placed on regenerative tissue-engineered and cell-based therapies that are expected to provide this missing part and either directly or indirectly instigate recovery of functional cardiac tissue.

For several decades, various cell-based therapies have been evaluated both preclinically and clinically but unfortunately, thus far, the approaches have had poor clinical translation due to high complexity, costs, lengthy preparation time, requirements for special production facilities, and sometimes frank incompatibility with clinical practice (5, 6). To add to the complexity, several cell sources and types, including stem and progenitor cells from embryonic tissue, adult mesenchymal or cardiomyocyte-differentiated stem cells, have been investigated (7). Many of these cell sources have been assessed in clinical trials, but none are used as a part of clinical practice (8).

The studies conducted in the early 2000s that addressed cardiac progenitor cells, first in the rodent and then in adult human heart (9, 10), generated considerable interest in functional regeneration of the heart. However, since the discovery of some major controversies concerning some of these studies, the concept of cardiac cell therapy has encountered ardent debates, emphasizing the need to re-evaluate the fundamental mechanisms underlying cardiac cell therapy. Nevertheless, cardiac progenitor cells have become a potential candidate for cell therapy (11, 12), and these cells may be superior to other cell types in terms of regenerative effects (13, 14). Although cardiac progenitor cells can be found in various locations throughout heart (12, 15-17), atrial tissues may contain the densest population (12, 18, 19). Moreover, cells isolated from atrial appendage tissues have demonstrated therapeutic potential (20). Our clinical AADC consortium recently demonstrated the clinical feasibility of direct intraoperative processing and epicardial transplantation of autologous atrial appendage micrografts (AAMs) (21, 22). Analysis of results from a more extensive clinical feasibility trial evaluating this approach in a limited number of patients is expected be reported later this year (ClinicalTrials.gov identifier: NCT02672163).

The aim of this study was to investigate the functional efficacy and molecular therapeutic mechanisms of epicardial AAM patch transplantation in a murine model of MI and HF. The overview of the study is presented in Figure 1.

**Figure 1.**
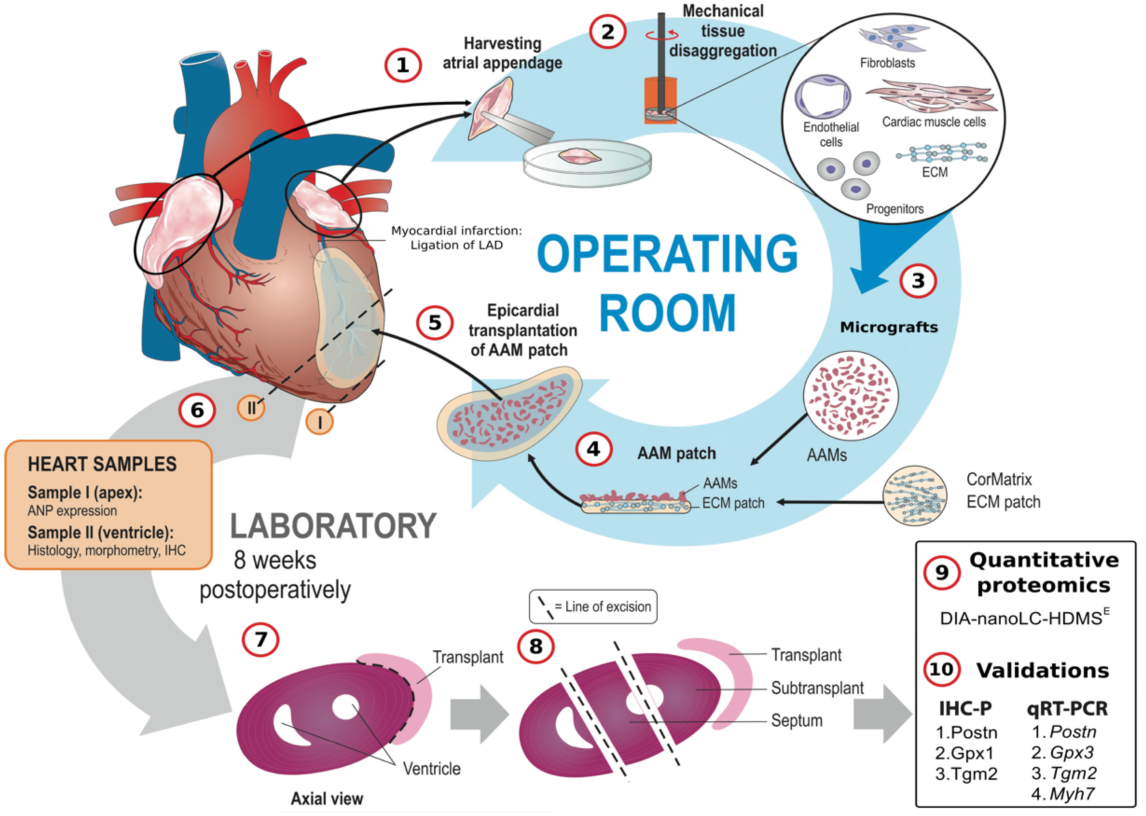
Overview of the study protocol. A schematic presentation of the processes carried out in the operating room and subsequently in the laboratory are illustrated. For further information, see the Detailed Materials and Methods. (**1**) For each AAM patch, both left and right atrial appendages were harvested from three syngeneic donor mice after placement of an atrial clip. (**2**) and (**3**) Harvested appendages were disaggregated to atrial appendage micrografts (AAMs) with a Rigenera machine. (**4**) An ECM sheet (CorMatrix® ECM™ for cardiac tissue repair) was peeled into four layers, and 1-ply sheets were cut into circles using an 8 mm diameter tissue punch. To create an AAM patch, we evenly dispersed the AAM suspension, containing tissue material from six appendages (three from both the right and left sides), onto the matrix sheet and sealed with a thin layer (10 μl volume) of fibrin tissue glue. (**5**) The AAM patch was subsequently fitted onto the recipient heart with the AAM-containing side facing the epicardium of the left ventricle at the site of infarction. The transplant was further fixed to the myocardium by three sutures to ensure that the patches remained in place. (**6**) After the 8-week follow-up, animals were sacrificed, hearts were excised, and ventricular tissue samples were collected for histology or snap frozen in liquid nitrogen. Apices were also collected for the measurement of ANP mRNA expression with qRT-PCR. (**7**) Thin histological sections were used for immunohistochemistry and morphometry. (**8**) and (**9**) Cryosections of 12-µm thickness were collected from three anatomic groups (transplant/patch, subtransplant and septum) for proteomic profiling. (**10**) Lastly, rest of the cryopreserved samples were used for selected validative qRT-PCR measurements, and paraffin embedded tissue samples for IHC-P staining.

## Results

### Myocardial function

Left ventricular ejection fraction (LVEF) with corresponding area under the curve (AUC) analyses spanning the 8-week follow-up period are presented in the left and right panels of Figure 2A, respectively. ECHO recordings demonstrated better functional recovery in the AAM patch group than in the ECM patch and MI groups. Lower initial LVEF values in the Sham group were due to lower weight of the group’s mice at the start of the experiment.

**Figure 2.**
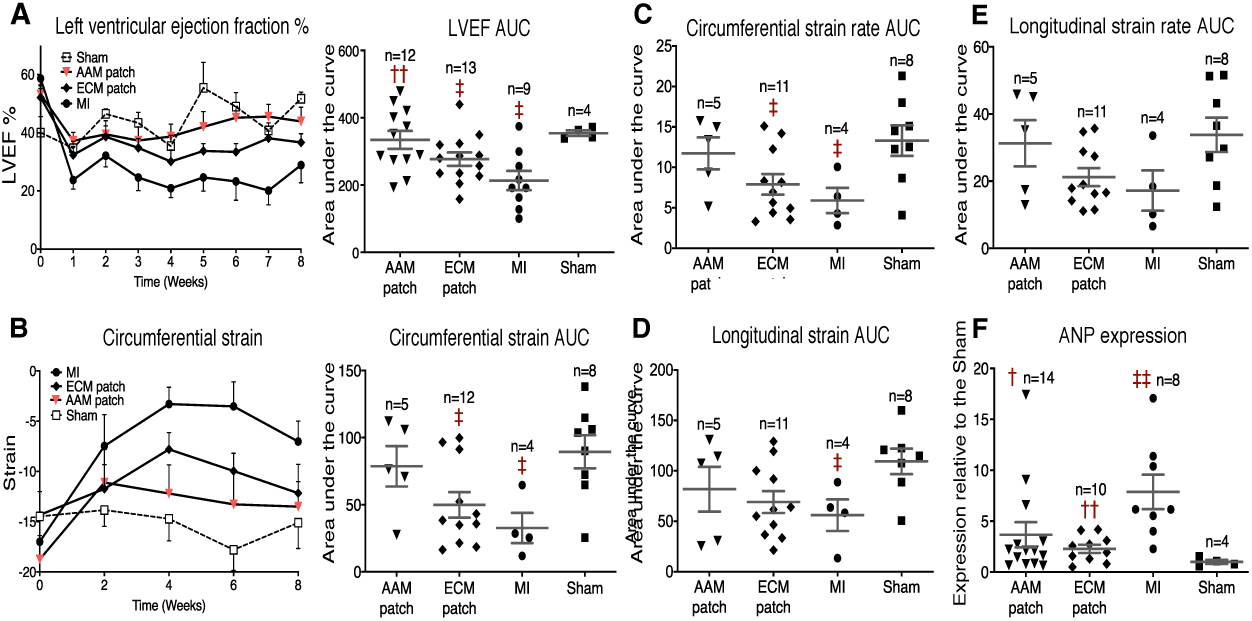
Myocardial function. (**A**) Left panel shows weekly performed LVEF measurements; right panel shows AUC analysis from the LVEF follow-up data. (**B**) Left panel shows the circumferential strain measured with ECHO at 2-week intervals; the right panel shows the corresponding AUC. (**C**) AUC analysis of the circumferential strain rate. (**D**) AUC analysis of the longitudinal strain; (**E**) AUC analysis of the longitudinal strain rate. (**F**) Relative expression of ANP mRNA to expression of 18S mRNA from apex tissue samples as analyzed by real-time PCR. Two-tailed Mann-Whitney *t*-test was applied and data normality assessed with Shapiro-Wilk test for normality. The scatter plot data represent mean ± SEM, while the n-values represent the number of mice measured. †/‡*P* < 0.05; ††/‡‡*P* < 0.01; †, *P* vs. the MI group; ‡, *P* vs. the Sham group.

In all groups undergoing intervention, a decline in LVEF was evident during the first two weeks of the follow-up. However, in the AAM and ECM patch groups, the fall in LVEF seemed considerably smaller in magnitude than that observed in the MI group. The acute declines in LVEF at one-week time-point postoperatively in AAM patch, ECM patch and MI groups were as follows: 18.03 (from 54.49±2.93% to 36.69±2.56%), 19.77 (from 52.15±3.10% to 32.38±2.96%) and 34.94 percent points (from 58.72±2.58% to 23.78±3.16%), respectively, with significant difference perceptible only between AAM patch and MI groups. However, since prevention of acute HF seemed evident in both patch groups, we hypothesize a considerable protective effect to occur by direct mechanical support, i.e. unloading, of the injured ventricle by the patch transplantation itself. The additional transplantation of micrografts might enhance the mechanical effect of the patch acutely, since the drop in LVEF was further reduced in the AAM patch group. In the later phase of the follow-up, a separation could be seen in the LVEF curves of the AAM and ECM patch groups, suggesting that the transplanted AAMs activated processes in the adjacent myocardium against the gradual development of chronic HF during a longer period of time. Significantly higher mean LVEF values were observed in the AAM group than in the MI group from 4^th^ week onward until the 7^th^ week of follow-up. At the end of follow-up, mean LVEF values reached 43.91±4.91% (AAM patch), 36.68±3.00% (ECM patch), 28.92±6.09% (MI) and 51.73±2.29% (Sham).

The LVEF measurements over the entire follow-up period were supplemented with AUC analysis (Figure 2A, right panel). The AAM patch group demonstrated highly preserved LVEF function based on AUC analysis, since significant decrease was observed only in ECM patch and MI groups compared against the Sham-level values. Furthermore, only the AAM patch group showed preservation of function when compared to MI values. The average LVEF AUC-values were 334.2±26.68 (AAM patch), 277.1±19.85 (ECM patch), 213.4±28.55 (MI), and 354±8.21 (Sham).

To provide further insights into the functional state of the left ventricle, we performed additional strain analyses during ECHO recordings. Comprehensive data from all strain analyses performed is presented in the online supplements (Figure S1 for the strain data, and Table S1 for all ECHO data) and summary of the AUC data is shown here in Figure 2B-E. Table 1 provides the mean values of ECHO parameters at the end of the follow-up. Overall, strain parameters in both patch groups were well preserved, with the greatest preservation in the AAM patch group, compared to the declines seen in the MI group. Summary of the strain parameter measurement results follows.

**Table 1.** Mean values of the ECHO follow-up at the 8-week time point. The data from the LVEF is presented more extensively in Figure 2A. */†/‡*P* < 0.05; **/††/‡‡*P* < 0.002; ***/†††/‡‡‡*P* < 0.0002; *, *P* vs. the matrix group; †, *P* vs. the MI group; ‡, *P* vs. the Sham group. Groupwise analysis was carried out using one-way ANOVA.

|  |  | AAM patch | ECM patch | MI | Sham |
| --- | --- | --- | --- | --- | --- |
| LVID (mm) |  |  |  |  |  |
|  | End diastolic | 4.82±0.22 | 4.93±0.16 | 5.07±0.19 | 4.22±0.11 |
|  | End systolic | 3.86±0.33 | 4.07±0.2 | 4.39±0.3 | 3.12±0.12 |
| IVS (mm) |  |  |  |  |  |
|  | End diastolic | 0.79±0.05 | 0.71±0.06 | 0.73±0.05 | 0.7±0.03 |
|  | End systolic | 1.13±0.09 | 0.98±0.09 | 0.97±0.09 | 0.99±0.06 |
| LVPW (mm) |  |  |  |  |  |
|  | End diastolic | 0.91±0.07 | 0.83±0.09 | 0.99±0.16 | 0.8±0.04 |
|  | End systolic | 1.13±0.09 | 0.96±0.1 | 1.08±0.16 | 1.02±0.01 |
| LVEF (%) |  | 43.91±4.91 | 36.68±3 | 28.92±6.09 | 51.73±2.29 |
| FS (%) |  | 22.46±2.95 | 17.88±1.66 | 14.31±3.6 | 26.25±1.42 |
| LV Vol (μL) |  |  |  |  |  |
|  | End diastolic | 111.8±11.61 | 116.3±9.01 | 124.1±9.84 | 80±5.12 |
|  | End systolic | 68.67±12.31 | 75.75±8.47 | 92.11±11.46 | 38.75±3.75 |
| SV (μL) |  | 43.17±1.62 † | 40.58±2.42 | 32±3.64 | 41.25±2.78 |
| HR (BPM) |  | 357.4±9 | 363.3±11.37 | 377.6±17.41 | 367.5±19.6 |
| CO (mL/min) |  | 15.38±0.64 | 14.82±1.07 | 11.89±1.25 | 15.28±1.8 |

Mean circumferential strain values (Figure 2B, left) at the 8-week time-point were - 13.5±2.48 (AAM patch), -12.2±2.87 (ECM patch), -7.0±2.04 (MI) and -15.1±2.57 (Sham). The supplementing AUC analysis (Figure 2B, right) revealed analogous results that were seen in LVEF AUC measurement with preserved function without significant decline only observable in the AAM patch group. The average AUC-values for circumferential strain were 78.6±15.04 (AAM patch), 49.9±9.50 (ECM patch), 32.6±11.27 (MI), and 89.46±12.30 (Sham). Mean circumferential strain rates at the end of the follow-up were -4.2±0.65 (AAM patch), -3.5±0.90 (ECM patch), -2.3±0.45 (MI) and -4.5±0.55 (Sham) and the corresponding AUC-values for the circumferential strain rate were as follows: 11.7±1.98 (AAM patch), 7.9±1.27 (ECM patch), 5.9±1.56 (MI), and 13.31±1.88 (Sham). Again, the same pattern of better recovery in the AAM patch group was demonstrable.

Alike with the circumferential parameters, longitudinal strain did not decline significantly in either patch group, and only the MI group against the Sham group showed significant decrease (Figure 2D). Longitudinal strain values at the 8-week time-point were - 13.4±2.16 (AAM patch), -16.6±2.83 (ECM patch), -12.9±4.05 (MI) and -14.4±2.41 (Sham) with the corresponding AUC-values being 81.8±22.23 (AAM patch), 69.1±10.87 (ECM patch), 56.1±15.65 (MI), and 109.4±12.67 (Sham). Further, the AUC analysis of the longitudinal strain rate did not present any statistically significant differences between groups (Figure 2E). Longitudinal strain rate values at the 8-week time-point were -5.0±0.63 (AAM patch), - 5.2±0.86 (ECM patch), -4.2±0.97 (MI), and -4.8±0.40 (Sham) with the corresponding AUC-values being 31.3±6.88 (AAM patch), 21.2±2.67 (ECM patch), 17.2±6.00 (MI), and 33.8±5.12 (Sham).

### Atrial natriuretic peptide expression

Myocardial samples from heart apices were analyzed for atrial natriuretic peptide (ANP) mRNA (Figure 2F) expression. ANP mRNA expression was significantly lesser in both patch groups (3.6±1.24 for AAM, and 2.3±0.41 for ECM) compared to level seen in the MI group (7.9±1.70). Interestingly, the AAM patch group demonstrated a trend for higher ANP production when compared to Sham (1.0±0.27) and ECM patch groups’ production rates. A plausible explanation for this elevated ANP mRNA expression could be that ANP was produced from the remnants of the transplanted micrografts, which we know to be atrial in origin with highly specialized capacity to produce ANP.

### Fibrosis

Build-up of fibrous tissue, analyzed by the scar transmurality index (i.e., scar thickness divided by the wall thickness at measurement point [%], Figure 3A) and ventricular wall thickness (Figure 3B) measures, was significantly attenuated in both patch groups. The mean scar transmurality index values were as follows: 50.2±3.69% (AAM patch), 53.1±3.54% (ECM patch), and 81.3±4.26% (MI) with the least amount of fibrotic material in the AAM patch group among the all interventional groups studied. The mean ventricular wall thicknesses were 0.21±0.01 cm (AAM patch), 0.16±0.08 cm (ECM patch), 0,061±0.071 cm (MI), and 0.15±0.094 cm (Sham). Both patch groups demonstrated preserved ventricular wall thickness compared to MI, while the increased thickness in the AAM patch group seems to be attributable to the transplanted micrografts. Planimetry analysis (Figure 3C) showed a lower degree of transmural infarction scar in both patch groups relative to the MI group. The angle of spread of collagenous scar tissue showed significant differences in both patch groups relative to the MI group. The exact mean angles of the infarction scar were 51.4±13.7° (AAM patch), 59.4±14.1° (ECM patch), and 152.4±9.0° (MI). Moreover, the mean angle of the epicardial fibrotic tissue represented 135.3±4.7° in the AAM patch group and 161.7±12.8° in the ECM patch group. Finally, representative histological images from each study group are shown in Figure 3D, with a clearly observable tissue level therapeutic effect following either AAM or ECM patch transplantation.

**Figure 3.**
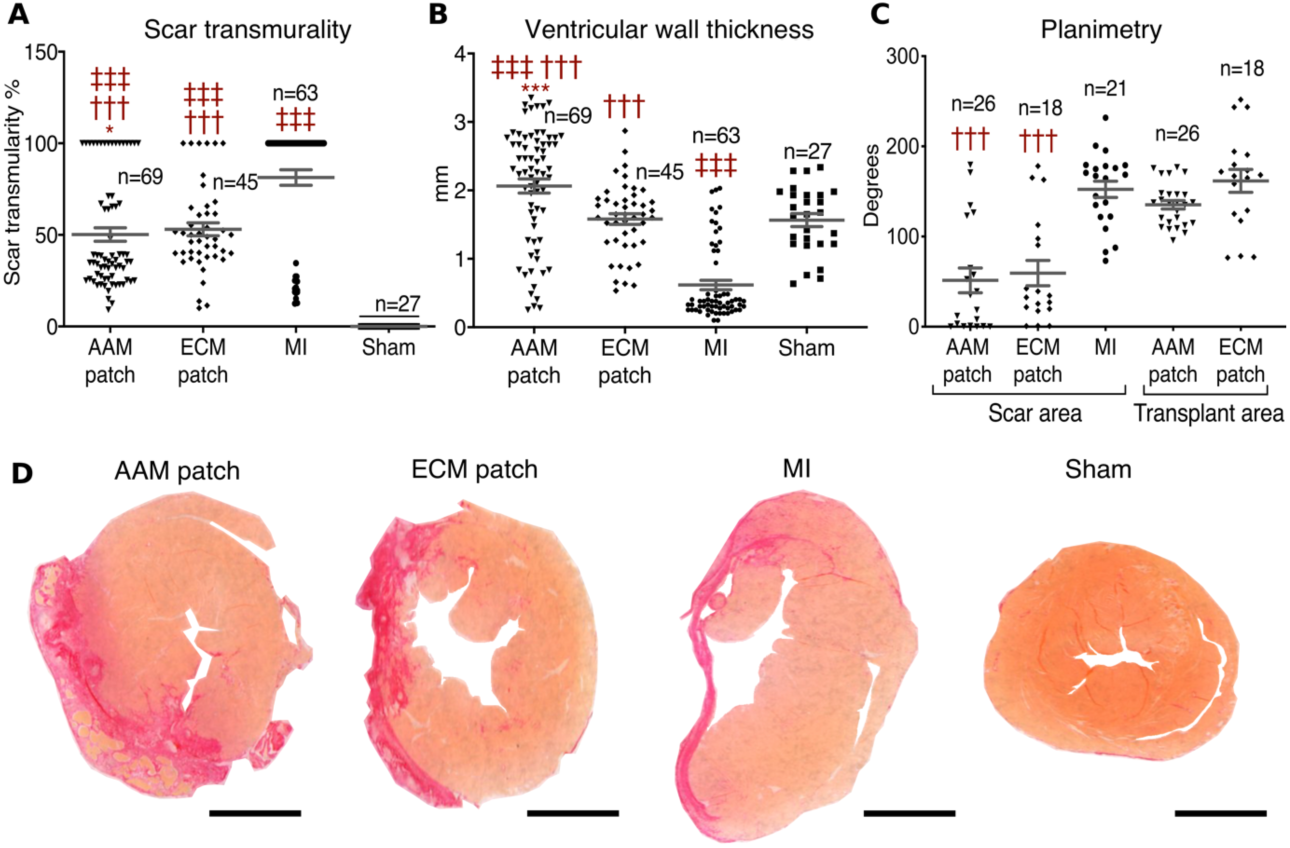
Ventricular morphometry. (**A**) Infarction scar transmurality was assessed by measuring the thickness of collagen-containing tissue and comparing it to the total thickness of the ventricular wall. (**B**) Ventricular wall thickness; thickness of the transplant was included in the measurement. (**C)** Planimetry analysis of the infarction scar angle. (**D**) Selected representative sections from the pool of Picrosirius Red-stained tissue sections from which the assessment of tissue morphometry and infarction scar transmurality was performed are presented. Scale bars equal 2 mm. Two-tailed Mann-Whitney *t*-test was applied and data normality assessed with Shapiro-Wilk test for normality. The scatter plot data represent mean ± SEM. The n-values represent the number of sites measured. */†/‡*P* < 0.05; **/††/‡‡*P* < 0.01; ***/†††/‡‡‡*P* < 0.001; *, *P* vs. the ECM patch group; †, *P* vs. the MI group; ‡, *P* vs. the Sham group.

In addition, the endocardial interstitial fibrosis was evaluated. Overall, the degree of tissue fibrosis in the endocardial surface was lower in both patch groups than in the MI group. The subendocardial surface was taken into focus, since this subregion in the heart is the least tolerant to hypoxia, and thus, the subendocardial surface is the first region to be damaged and scarred after an ischemic insult. Evaluation of the endocardial fibrosis, with collagen subtype characterization, is presented in Figure 4. In all analyses (total, green, orange and bright fibers), both patch groups demonstrated significantly lesser total collagen content when compared to the MI group. The mean content of collagen fibers represented 3.23±0.83% (AAM patch), 1.67±0.55% (ECM patch), and 15.2±1.96% (MI); second, for bright fibers 0.035±0.013% (AAM patch), 0.005±0.002% (ECM patch), and 0.309±0.088% (MI); third, for green fibers 0.77±0.19% (AAM patch), 0.28±0.062% (ECM patch), and 4.54±0.65% (MI); fourth, for orange fibers 2.43±0.65% (AAM patch), 1.38±0.53% (ECM patch), and 10.32±1.32% (MI).

**Figure 4.**
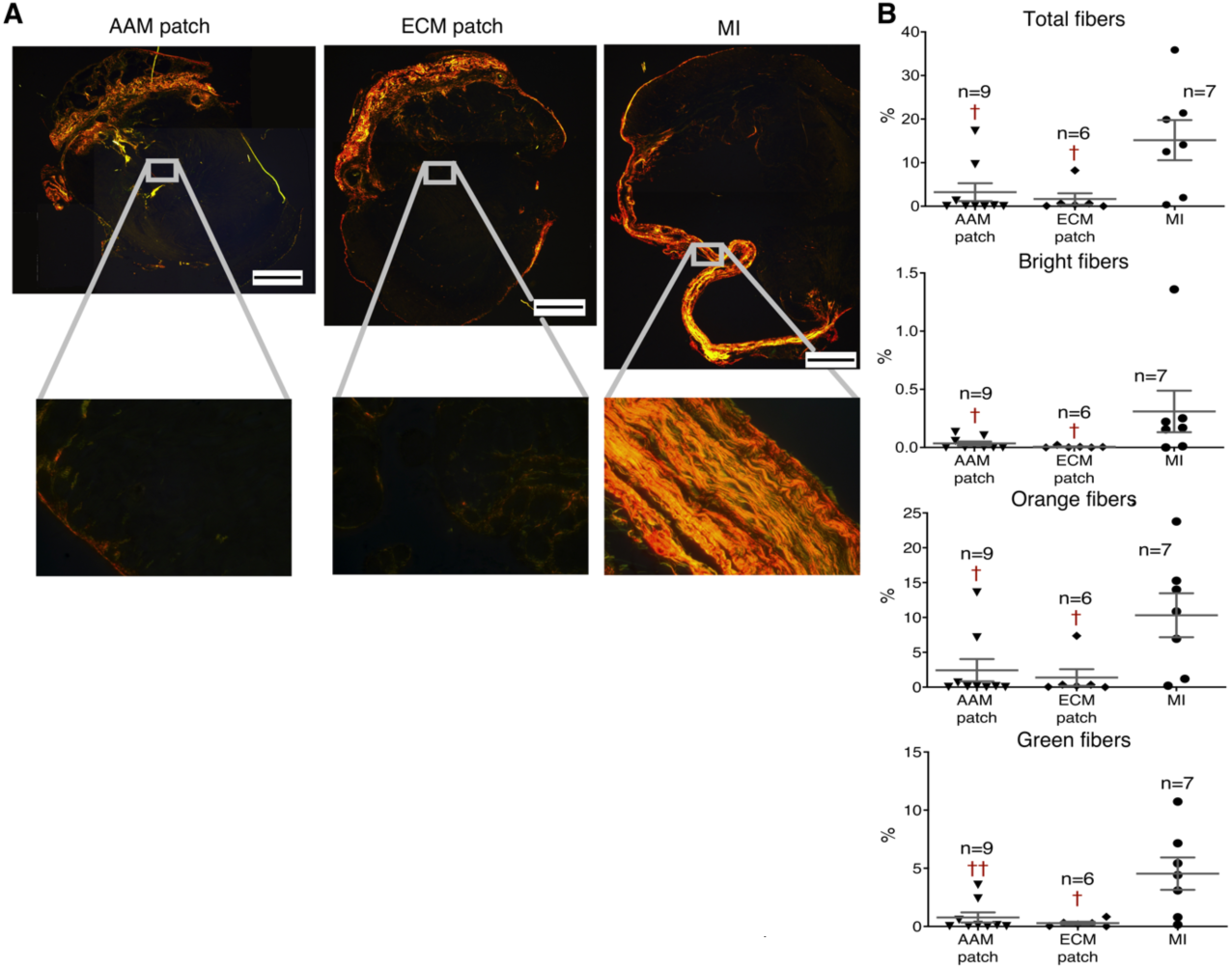
Evaluation of fibrosis. (**A**) Illustration of collagen analysis. Upper pictures are representative illustrations of the cardiac tissue sections photographed with circularly polarized light. Lower magnified and representative images were obtained from the endocardial surface of the infarction area from the left ventricle using 40 x magnification. The scale bar equals 1 mm. (**B**) Quantitative analysis of collagen fiber subpopulations from the endocardium. The uppermost panel shows the total content of Picrosirius Red-positive fibers in the tissue samples analysed, while the panels below present the amount of the birefringent, orange and green fibers seen with circularly polarized light, respectively below. Using Picrosirius Red staining and circularly polarized light, thinner type-III fibers appear green, whereas thicker and more mature type-I fibers appear red or yellow. Two-tailed Mann-Whitney *t*-test was applied and data normality assessed with Shapiro-Wilk test for normality. The scatter plot data represent mean ± SEM, while the n-values represent pooled means (7 measurement sites) from each individual sample measured. †*P* < 0.05; ††*P* < 0.01; †, *P* vs. the MI group.

### Histological preservation of myocardium

We evaluated the preservation of myocardial tissue with anti-cardiac troponin T (cTnT) immunohistochemistry (Figure 5A-B) and by quantifying the area of unstained tissue from sections stained with Picrosirius Red for collagen (Figure 5C-D). The area of tissue stained positive for cTnT in the subtransplant tissue seemed to be larger in both patch groups when compared to the MI group. However, clear tissue-level damage from the artificial LAD ligation was observable across all interventional groups, since significant reduction against the Sham was demonstrable in all interventional groups. Using the cTnT positivity in the Sham group as a reference, cTnT positive tissue areas in the subtransplant encompassed 49.4±1.94% (for AAM patch), 53.6±2.46% (ECM patch), 35.0±2.28% (MI), and 99.68±3.52% (Sham).

**Figure 5.**
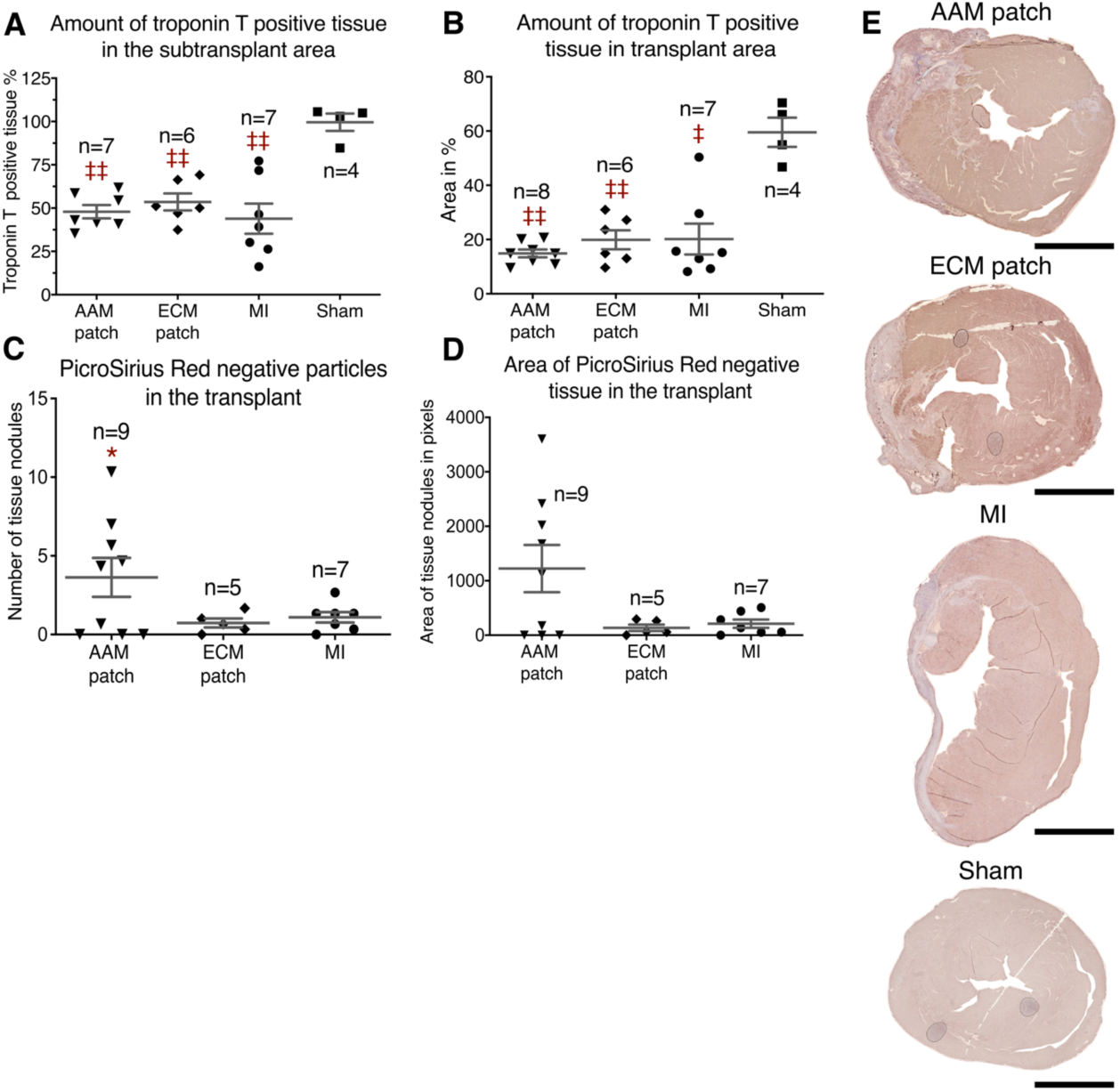
Histological preservation of myocardium. (**A**) Quantitation of Troponin T positive tissue content in the adjacent myocardium to the transplant, the subtransplant zone. (**B**) Similarly, quantitation of anti-troponin-T positively stained area in the transplant. (**C**) Number of nonfibrous tissue nodules larger than 100 pixels inside the infarction scar (or the transplant area in AAM and ECM patch groups) in the Picrosirius Red-stained samples. (**D**) Area of the nonfibrous, Picrosirius Red-negative tissue nodules. MI group values are presented as a reference. (**E**) Selected representative sections from the immunohistochemical staining set with anti-troponin T antibody. The scale bar equals 2 mm. Two-tailed Mann-Whitney *t*-test was applied and data normality assessed with Shapiro-Wilk test for normality. Scatter plot data represent mean ± SEM, while the n-values represent pooled means (9-29 measurements) from each individual sample measured.*/‡*P* < 0.05; **/‡‡, *P* < 0.01; *, *P* vs. the ECM patch group; ‡, *P* vs. the Sham group.

Subsequently, the amount of nonfibrous tissue inside the infarction scar and transplant/patch was analyzed. Figure 5C shows the quantity of tissue nodules within the site of infarction. Only tissues that did not stain with Picrosirius Red were measured. Significantly more nodules were found in the AAM patch group than in both the ECM patch and MI groups, possibly suggesting a preservation of transplanted AAMs. The mean counts of noncollagenous tissue nodules were 3.6±0.71 (AAM patch), 0.7±0.28 (ECM patch) and 1.1±0.24 (MI). The nodules observed in the AAM patch group were also the largest, and this difference was significant compared to those in the ECM patch and MI groups (Figure 5C). The mean Picrosirius Red-negative tissue area encompassed 1205±258 pixels (AAM patch), 141±49 pixels (ECM patch), and 258±75 pixels (MI) (see Figure 5D). However, after subsequent analyses for cTnT staining, these nodules remained cTnT negative (Figure 5B), suggesting that transplanted AAMs do not contribute directly to contractile cardiac tissue. The mean relative cTnT positive tissue area was 15.1±0.55% (AAM patch) and 18.1±1.15% (for ECM patch). The reference value of cTnT content in the infarction scar was 15.2±1.35% (MI group), and the mean relative amount of cTnT-stained tissue of the ventricular wall in the Sham group was 58.6±2.34%. Figure 5E presents representative histological sections from each study group stained against cTnT.

### Proteomics

In a separate experiment designed to compare AAM and ECM patch molecular tissue-level responses utilizing the same MI model and the 8-week follow-up as previously described, label-free site-targeted proteomics was performed on samples from AAM and ECM patch groups and from the following areas: (1) ECM or AAM patch (transplant), (2) subtransplant and (3) distal septal myocardium (see Figure 1). Our aim was to elucidate the mechanisms accounting or responsible for preserved cardiac function and decreased scar formation demonstrated by preserved ECHO follow-up parameters, collagen analysis, Picrosirius Red and cTnT stainings. All quantified and differentially expressed proteins (DEPs) are presented in Data File S1 and the proteomics data have been uploaded to the Mass Spectrometry Interactive Virtual Environment (MassIVE) for in-depth review (see Supplemental data). The quantified proteins (1,005 in total) were assessed utilizing a Spearman correlation heatmap, which confirmed site-specific dissection of the transplant, subtransplant and remote septal samples. Samples from the patch transplant area clustered as a separate group, while the myocardial samples from the subtransplant and septal areas formed another major cluster. Within this myocardial cluster, samples from the AAM patch-treated group from both subtransplant and septal areas clustered separately from ECM patch samples without micrograft therapy, suggesting a widespread therapeutic effect of AAMs (see Figure S2). By comparing the AAM patch group to the ECM patch control group, we identified 293 DEPs (45 upregulated and 248 downregulated) in the transplant area. In the subtransplant area, 216 DEPs were characterized (151 upregulated and 65 downregulated), while in the septal area, 43 DEPs were identified (42 upregulated and 1 downregulated). All DEPs were scrutinized using Ingenuity^®^ Pathway Analysis (IPA^®^) software to reveal associated canonical pathways, biological functions and disease states. Overview of the results are presented next.

Transplant/patch zone—DEPs associated with the pathways for *Oxidative phosphorylation* (activation), *Actin cytoskeleton signaling* (activation), *Sirtuin and calcium signaling* (inactivation) were altered in the AAM patch transplant zone compared to those in the ECM patch zone without AAMs (Figure 6A). Utilizing the diseases and functions feature, we detected elevated processes related to the survival of transplanted cells in the AAM patch. Specifically, functional categories including *Cell death and necrosis* were significantly downregulated (B-H z-scores -2.99 and -3.43, respectively; Figure 6B), while those related to *Cell survival and cell viability* were upregulated (+3.09 and +3.36, respectively; Figure 6B).

**Figure 6.**
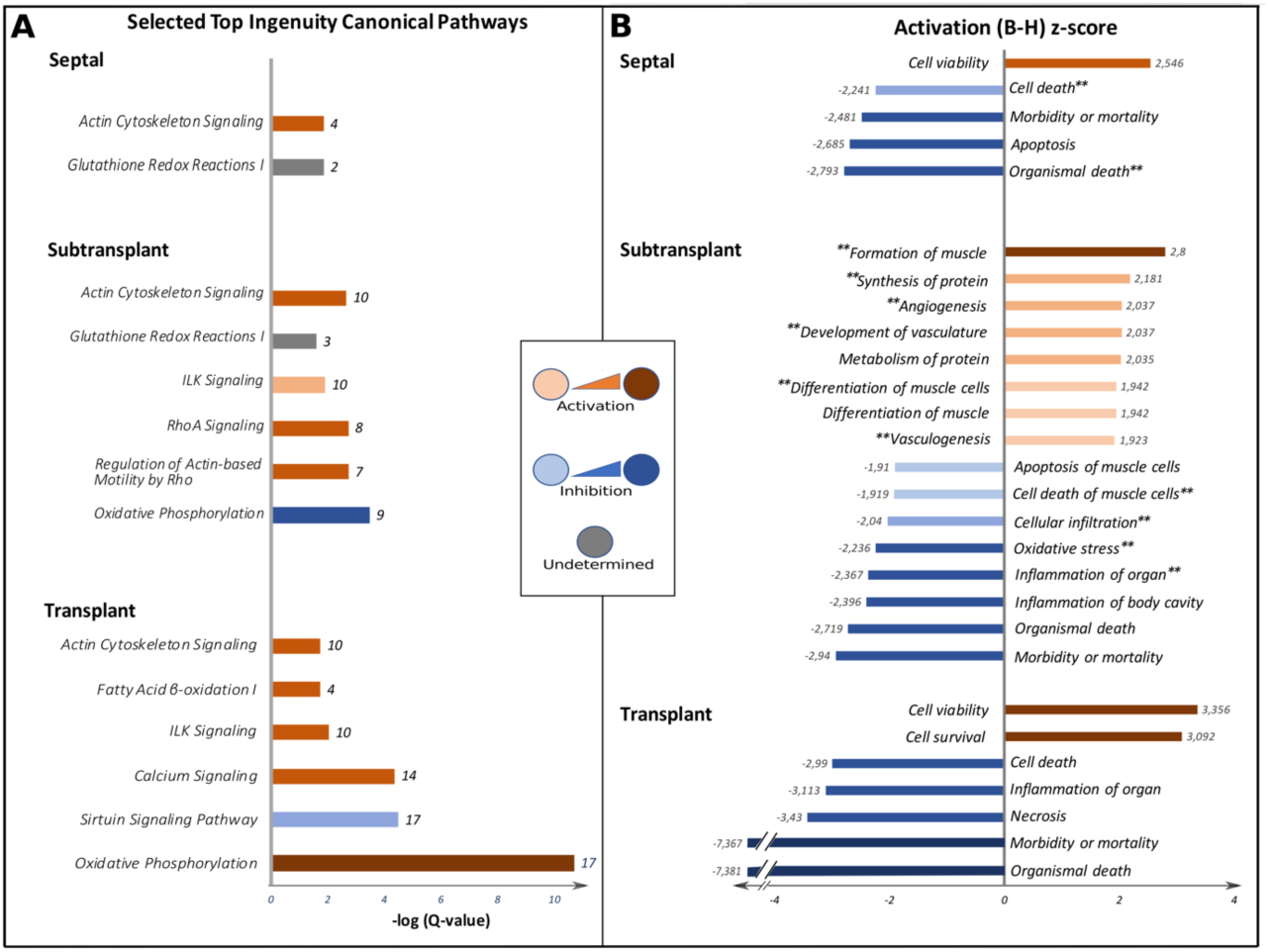
Proteomics, canonical pathways and biological functions. From each anatomical site, selected canonical pathways (**A**) and functions (**B**) are shown. The results are based on IPA® software analysis of given representative locational DEPs (shown in Data File S1). The Q-value represents the B-H *P*-value (Benjamini-Hochberg corrected *P*-value). Q-value ≥ 1.5 and |z-score| ≥ 2 were used as criteria for significant activation or inactivation of a given pathway or process. All detected changes were compared to the ECM patch group, thus, these changes were attributed to AAMs transplantation. At the end of each bar in the subplot (**A**) is the number of DEPs associated with a given pathway while in (**B**) the approximate z-score values are presented. Functions annotated with double asterisk (**) are shown further in Figure 7. All canonical pathways as well as diseases and functions identified in each anatomical site are provided in Figures S3 and S4, respectively.

Subtransplant zone—Using the same features used for the analysis of the transplant/patch zone, pathways related to cytoskeletal changes and reorganization were activated in the AAM patch-treated subtransplant zone, namely, *Regulation of actin-based motility by Rho, RhoA signaling* and *Actin cytoskeleton signaling* (Figure 6A). From a metabolic perspective, *Oxidative phosphorylation* was predicted to be significantly decreased, whereas *Glutathione redox reactions I* were significantly altered with increased expression of enzymes GPX3, GPX1, and GSTM2 in the AAM patch-treated group (Figure 6A and Data File S1). Notably, functions related to muscle cells’, especially cardiomyocytes’, viability and new muscle formation (*Formation of muscle* [B-H z-score +2.80], *Differentiation of muscle and muscle cells* [+1.90], *Apoptosis and cell death of muscle cells* [-1.91, -1.92, respectively]) were predicted to be significantly activated (Figure 6B). The inflammatory response (*Inflammation of organ* [-2.367]) and *Oxidative stress* (−2.236) showed decreased activity in the AAM patch-treated group compared to those in the ECM patch group (Figure 6B). Interestingly, we found a 1.518-fold increase in expression level of matricellular protein periostin (POSTN) and 1.663-fold overexpression of tissue transglutaminase-2 enzyme (TGM2) with AAMs transplantation (Figure 7 and Data File S1). Together, these changes provide a molecular-level insight into the observed functional improvements.

**Figure 7.**
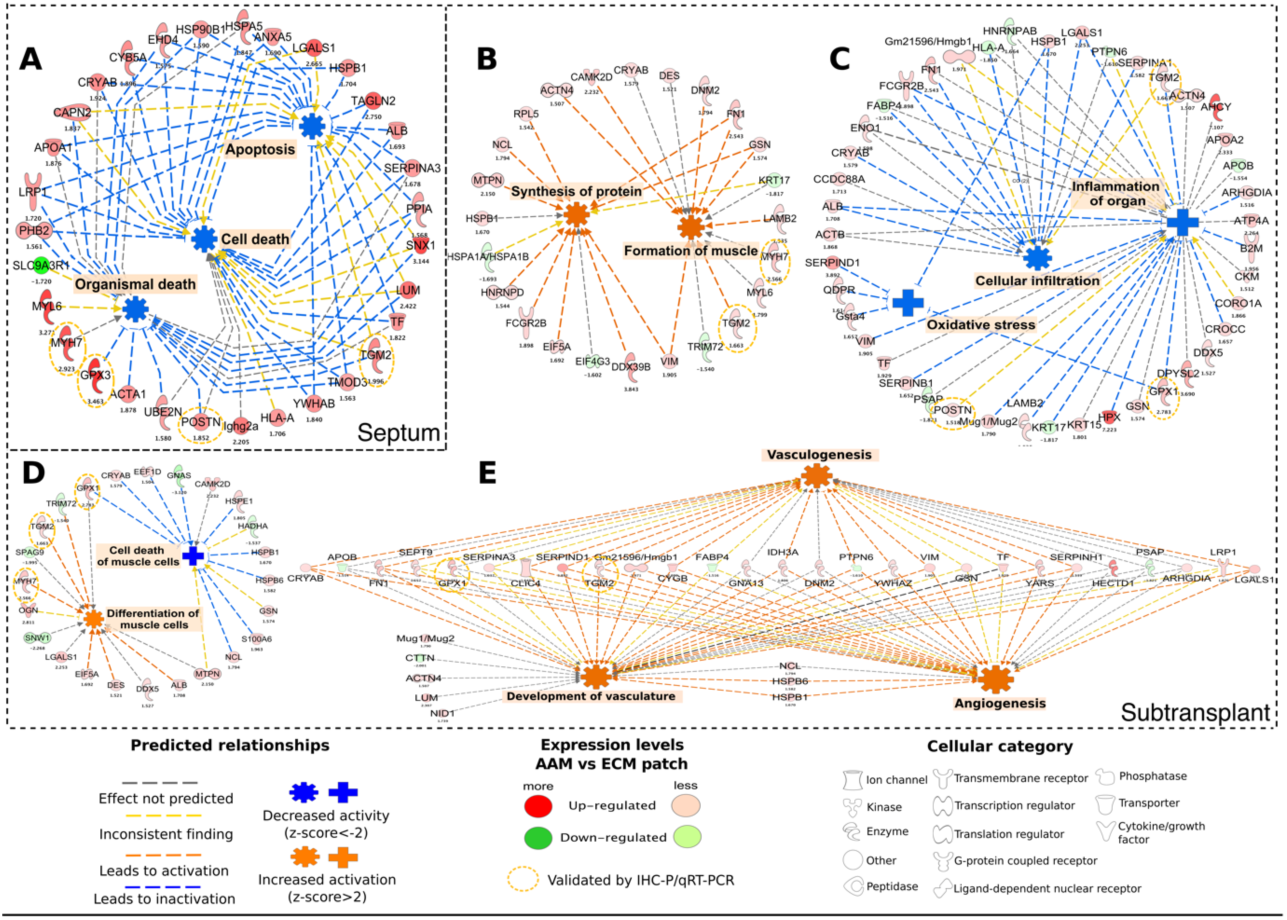
DEPs and associated biological functions. Selected biological functions from Figure 6 and associated DEPs are shown here. (**A**) Prosurvival functions and associated DEPs from the remote septum are presented. Periostin (POSTN) is documented cardiomyocyte mitogen and repair-mediator after ischemic cardiac damage. (**B**) Functions related to anabolic protein and muscle metabolism from the subtransplant are shown. Desmin (DES) is a pivotal intermediate filament in cardiomyocytes while Myosin 7 (MYH7) is a major slow twitch myosin isoform in the heart. (**C**) DEPs associated with a decreased inflammatory reaction in the subtransplant are shown. Glutathione Peroxidase 1 (GPX1) is a ubiquitous enzyme with antioxidant properties catalysing the hydrolysis of hydrogen peroxide radical to water. (**D**) DEPs associated with increased cardiomyocyte differentiation in the subtransplant immediately next to AAM patch transplantation are shown. However, these functions showed only substantially strong trends towards activation and inactivation with a |z-score| = 1.9 for both functions (with criterion for significant activation or inactivation being |z-score| ≥ 2.0). (**E**) Functions related to angiogenesis with associated DEPs in the subtransplant are shown. Transglutaminase 2 (TGM2) catalyses crosslinking of proteins and is associated in apoptosis, it is also documented to be prosurvival factor for cardiomyocytes following ischemic insult. Small numeric counts below each DEP represent the expression ratio in AAM patch group divided by that of the ECM patch group. Statistical significance for differential protein expression was assessed using two-way ANOVA with *P* criterion ≤ 0.05.

Septal zone—Similar to the subtransplant area, *Actin cytoskeleton signaling* and *RhoA signaling* was predicted to be activated. *Glutathione redox reactions I*, with increased expression levels of GPX3 and GSTM2 enzymes, was also demonstrated to be significantly altered (Figure 6A, Data File S1). Functions related to more diffuse prosurvival effects in remote septum seemed increased at the cellular, organ, and whole-organism levels in the AAM patch-treated group. At the cellular level, *Cell viability* was predicted to be activated (z-score +2.546), while *Cell death* and *Apoptosis* were decreased (z-scores -2.241 and -2.685, respectively, Figure 6B). At the system level, the functions *Morbidity and mortality* and *Organismal death* were predicted to be decreased (z-scores -2.481 and -2.793, respectively, Figure 6B). Notably, overexpression of both matricellular protein POSTN (1.852-fold) and TGM2 enzyme (1.996-fold) were also detected following AAM transplantation, far away from the initial AAM patch transplantation site (Figure 7).

While selected canonical pathways are presented in Figure 6, all associated canonical pathways as well as biological functions and disease states are presented in Figures S3 and S4, respectively. Selected biological functions and disease states are shown in Figure 7 from septal and subtransplant sites with associated DEPs and their expression level ratios calculated against the expression levels in ECM patch-treated controls. Supplemental Table S2 further presents all associated DEPs to each canonical pathway, biological function and disease state presented in Figure 6.

### Immunohistochemistry and qRT-PCR

To qualitatively assess the expression of selected differentially expressed proteins identified in various functional categories, we performed immunohistochemical staining for POSTN, TGM2 and GPX1 (see Figure 8A). POSTN expression localized specifically to the extracellular space of the scar-myocardium interphase mostly in the subtransplant beneath the patch. This kind of spatial staining pattern is well in line with the earlier reports on POSTN expression in the heart after acute MI (23). On the other hand, TGM2 expression mainly localized to the patch and intracellular compartment of the cardiomyocytes near the minor scar areas while GPX1 expression demonstrated less pronounced staining but localized in the myocardium to the cardiomyocytes near the scar areas and vascular walls and more intense in the patch areas. Results of qRT-PCR confirmed overexpression, in both subtransplant and septal zones, for *Postn, Myh7* and *Gpx3* (Figure 8B). The mRNA expression levels for *Postn, Myh7* and *Gpx3* were significantly higher only in AAM patch group when compared to the Sham (*P*=0.010, *P*=0.014 and *P*=0.028 respectively). Moreover, the expression of *Gpx3* was higher in AAM patch compared to ECM patch group (*P*=0.046). *Tgm2* mRNA levels did not show significant changes between groups. When comparing the results from gene and protein expression, it is important to consider the different biological regulation and temporal dynamics at the level of gene transcription and mRNA translation, since it can vary substantially (24).

**Figure 8.**
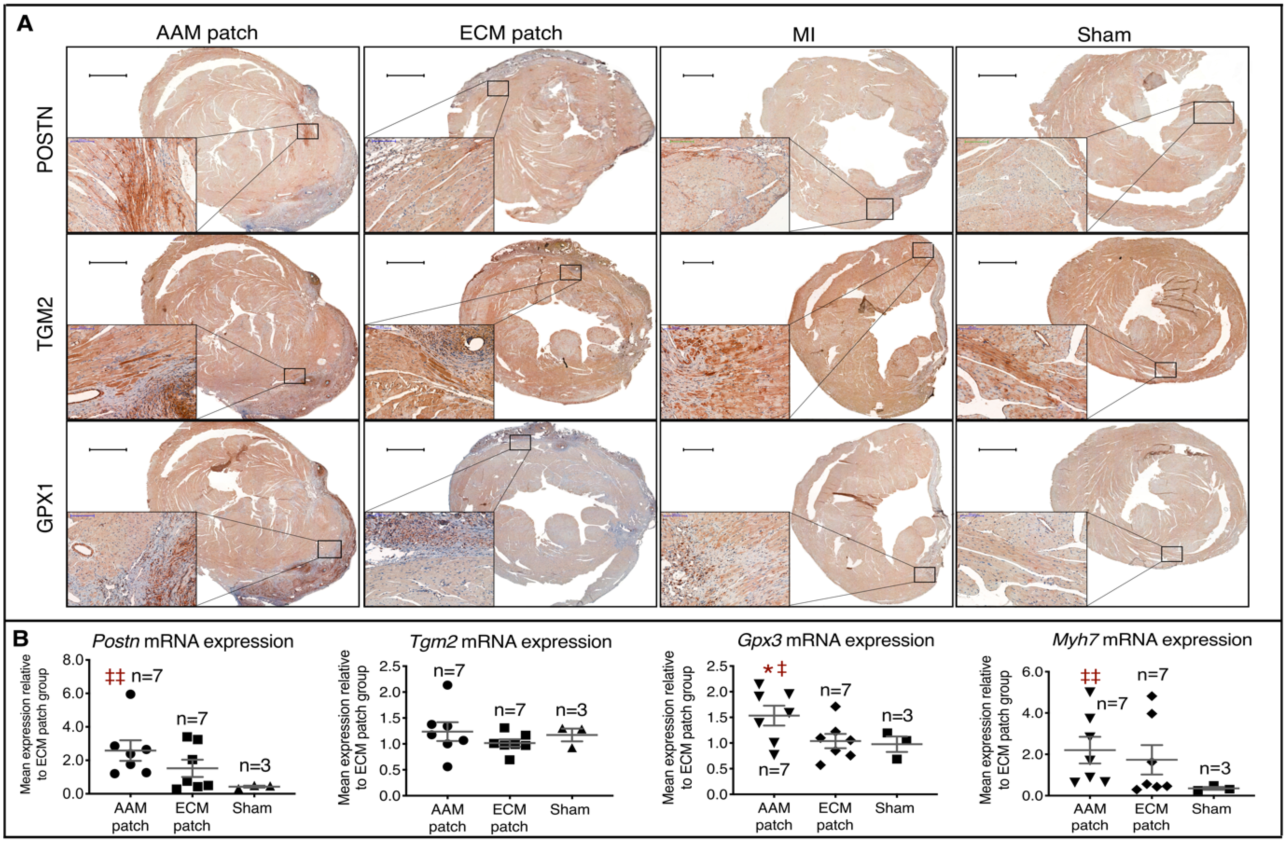
Validative IHC-P and qRT-PCR analyses. The set of samples preserved in paraffin was used for evaluating the fibrotic content and overall histomorphology of the study groups by IHC-P. Assessment by qRT-PCR was applied for the same set of cryopreserved samples as in the quantitative proteomics. (**A**) The representative images with magnifying subcaptions from the IHC-P stainings are presented here for three selected proteins of interest: POSTN, TGM2 and GPX1 (n=4 for each). POSTN exhibited marked expression in the patch zone as well as in the extracellular interphase of fibrotic scar areas and myocardium in all groups. TGM2 expression was mainly localized to the intracellular compartment of cardiomyocytes and endothelial cells, with the most pronounced expression localizing to the cardiomyocytes with immediate contact to the fibrotic areas in the subtransplant zone right below the patch. Finally, the GPX1 exhibited marked expression in the patch itself and in the vascular wall. Intracellular expression in cardiomyocytes seemed to be more pronounced in AAMs group as compared to the ECM group. The black, green and blue scale bars equal 1 mm, 200 μm and 100 μm, respectively. (**B**) The results of the qRT-PCR. Transcription of *Myh7, Gpx3* and *Postn*, together with the production of the corresponding mRNAs, was significantly higher after AAM patch transplantation (n=7) in comparison to the ECM patch transplantation (n=7), when both were compared against the steady state gene expression represented by the Sham expression levels (n=4).*Tgm2* expression was not altered. The statistical tests were calculated using one-way unpaired *t*-test with Welch’s correction for unequal variances. Normality of the data was assessed with Shapiro-Wilk normality test. The scatter plot data represents mean ± SEM, while the n-values represent the individual samples measured. */ ‡*P* < 0.05; **/‡‡*P* < 0.01; *, *P* vs. the ECM patch group; ‡, *P* vs. the Sham group.

## Discussion

We investigated the effects and tissue responses of epicardially transplanted ECM and AAM patches in a murine model of MI and HF. AAM patch therapy demonstrated significantly improved cardiac function by ECHO. Treatment with either ECM or AAM patches greatly preserved myocardial structure, attenuated ventricular wall scarring, and dramatically restricted tissue damage. Myocardial site-targeted proteomics revealed the cell and molecular-level effects of AAMs spanning tissues under the transplant, reaching the myocardium. AAM therapy was associated with activated pathways for cardiomyogenesis, angiogenesis and anti-inflammation, as well as reduced oxidative stress.

Cell-based therapies and tissue engineering have long been investigated for their ability to deliver tissue regenerative adjuvant support. Atrial appendages were identified as niches for cardiac stem and progenitor cells and serve as a good autologous source of myocardial tissue for therapy due to their easy accessibility and relative redundancy for cardiac function (18, 19). Atrial appendage tissue is often removed and discarded during cardiac surgery to prevent formation of thrombi and predisposing the patient to thromboemboli.

In our approach, atrial appendage tissue was mechanically disaggregated after harvesting to yield micrografts (25). An AAM patch was generated by combining the micrografts in a small amount of fibrin with a tissue-engineered ECM patch that was epicardially transplanted onto the ischemic myocardium. We recently reported a successful clinical application of the AAM patch during coronary artery bypass graft surgery (21,22). In cardiac MRI and by clinical evaluation 6 months after operation, the treatment demonstrated favorable safety and feasibility profiles. This was succeeded by an open clinical feasibility trial, the results of which are expected to be reported later this year (ClinicalTrials.gov identifier: NCT02672163).

Here, we compared the effects of micrograft-containing AAM patches to those of corresponding acellular ECM patches in a mouse model of ischemic HF. Our results demonstrated myocardial tissue protection, attenuated scarring, and retained cardiac function after epicardial transplantation of AAM patches. These effects were associated with micrograft persistence within the transplant, suggesting that these cellular beacons emit therapeutic signals in a paracrine manner to the myocardium, a recent and emerging new hypothesis for cellular adjuvant therapies for cardiac regeneration (26, 27).

We utilized mass-spectrometry-based quantitative proteomics to decipher those myocardial responses specific to AAMs by comparing tissue responses to matrix-patch transplants with or without AAMs after MI. We performed myocardial site-specific collection of samples from transplants, as well as from subtransplant and remote septal myocardium. Our results demonstrate a widespread cardioprotective effect on the tissue site under the transplant that also reached remote myocardial tissues. We discovered significantly decreased oxidative metabolism and oxidative stress and increased glutathione metabolism in the subtransplant area. These changes suggest an AAM-induced change in myocardial metabolism towards glycolysis, a process associated with an increased regenerative capacity of mouse myocardium in multiple studies (28-32). Thus, we propose that transplanted atrial micrografts induced, likely through a paracrine mechanism, a metabolic shift from oxidative to glycolytic metabolism in the ischemically challenged subtransplant area and a pro-reparative microenvironment for surviving cardiomyocytes, ultimately setting the stage for functional recovery. This effect is further supported by our observation that functions such as “*Formation of muscle*” and “*Differentiation of muscle cells*” were activated. Moreover, restoration of functional myocardial tissue is not merely a process of cardiomyocytes, but necessitates formation of new ECM, blood vessels, and lymphatics, which are mostly orchestrated by cell types other than cardiomyocytes. Our findings demonstrate increased angiogenesis, vasculogenesis, and widespread prosurvival changes in the myocardium, which indicate induction of a protective and restorative myocardial environment by AAMs. Interestingly, our results from proteomics showed significantly increased expression levels of the cardiomyocyte mitogen and matricellular protein POSTN (33, 34) in both the subtransplant and remote septal zones, thus further strengthening the cellular and molecular mechanisms behind the observed functional improvement after AAM patch transplantation.

The secreted matricellular protein POSTN mediates cell-matrix interactions and repair after tissue injury. As shown earlier, our data support the association of POSTN expression and myocardial regenerative signaling via the Rho-kinase pathway (23). During tissue repair and healing after myocardial infarction, POSTN has been shown to be secreted to cardiac interstitium mainly by activated cardiac fibroblasts (23). POSTN can stimulate cardiomyocyte re-entry to the cell cycle and prevent myocardial rupture after infarction (23, 33). We demonstrated here that epicardial AAMs patch therapy resulted in myocardial salvage and functional restoration and was associated with increased periostin gene and protein expression in subtransplant and septal myocardium. We propose that the AAMs-patch-induced expression of periostin taken together with enhanced neoangiogenesis and anti-inflammatory response promotes formation of a pro-reparative myocardial microenvironment.

In the subtransplant myocardium, AAM patch transplantation increased myocardial expression of proteins associated with muscle formation, vasculature development, and glutathione metabolism compared with those in infarcted myocardium treated with ECM patch without micrografts. DEPs were also negatively associated with cellular infiltration. In both subtransplant and septal myocardium, remote to the transplant site, we discovered increased expression of proteins associated with cardiomyocyte contraction (including MYH7, MYL1, MYL6, MYL12B, ACTA1, ACTN4, DES and CAMK2D). Interestingly, the expression of proteins involved in glutathione metabolism and antioxidant defenses (GSTM2, GSTA4, GPX3) were also significantly increased in the subtransplant and remote septal sites treated by AAM patch therapy compared with those treated by ECM patch therapy. The observed *Gpx3* gene expression difference between AAM and ECM patch groups further strengthens this association of the AAMs mechanism of action via enhanced myocardial antioxidant defenses. In addition, we found a significant increase in cell viability and a significant decrease in cell death and apoptosis in the septal zone in AAM patch-treated hearts. This prosurvival action is a documented paracrine effect of cardiac stem/progenitor cell therapy (35).

Some concerns have been raised about potential inflammation and foreign-body effects caused by the transplantation of xenogeneic material, but a systematic review advocates the safety of CorMatrix (ECM patch) in human cardiac surgery (36). MI, cell damage, and necrotic death of cardiac cells initiate an intense inflammatory response. Although the main purpose of inflammation is the removal of cellular debris and nonfunctional cells, the immunomodulatory actions exerted by infiltrating cells may contribute to tissue repair (37, 38). M1 versus M2 macrophages are attributed with differential roles in the modulation of the inflammatory response. M1 macrophages are associated with increased cardiomyocyte injury, whereas M2 and endogenous cardiac macrophages are associated with more rapid degradation of dead cells and a suppressed immunological response (39-41). We detected a decreased inflammatory reaction in the subtransplant zone challenged with ischemia and near the AAMs. We also identified increased angiogenesis and a strong trend towards increased muscle cell differentiation in this same zone with AAM patch transplantation. Thus, our results indicate that AAM patch transplantation might be modulating the inflammatory response following MI in a way that promotes functional myocardial recovery. However, further targeted studies on the inflammation-modulating mechanisms of epicardial AAM patch transplantation are warranted.

In addition to the function of transplanted atrial appendage micrografts, the ECM patch itself likely possesses cardioprotective effects. This assumption is supported by several observations. First, LVEF, one of the most widely used parameter to assess cardiac pump function in clinical settings, remained more than 10 percentage points higher in both the ECM and AAM patch-treated groups acutely at the one-week time point postoperatively (Figure 2A), demonstrating the profunctional effect of the ECM patch transplantation itself onto the ischemic myocardium. Second, the same phenomenon with other functional ECHO follow-up parameters such as circumferential strain and strain rate was detected (Figure 2). However, the AAM patch-treated group showed a trend towards improved LVEF during the longer 8-week follow-up period when compared to the ECM patch group; we propose that this phenomenon is attributable to AAMs transplantation and is most probably transmitted through the previously identified and discussed molecular and cellular processes. Third, the transmurality index and ventricular wall thickness measures of the infarction scar site (Figure 3) showed significant improvements with ECM patch transplantation alone at the 8-week time point postoperatively. Here, although the AAM patch group pinpointed still further enhancement in the functional parameters from the ECM patch group, the effect between ECM patch and MI group is quite notable. These findings underline the crucial role of the ECM patch transplantation itself.

As a supportive structure, an epicardial ECM patch can alter the biomechanics of the failing left ventricle in a beneficial manner. The total work done by the heart (***W***) during one cardiac cycle (systole+diastole) is the sum of (a) pressure-volume work (*PV*, “the context where the heart has to work”), (b) kinetic energy (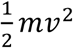, “the momentum that heart imparts to the blood”) and (c) tension heat (*kTt*, “the heart’s endogenous energy consumption”):

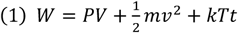

***P***, luminal pressure; ***V***, stroke volume; ***m***, mass; ***v***, velocity; ***k***, proportionality constant; ***T***, wall tension; ***t***, elapsed time

Wall tension (***T***) in the equation (1) is calculated as:

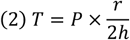

***P***, luminal pressure; ***r***, luminal radius; ***h***, myocardial wall thickness

Tension heat represents approximately 90-95% of all work done by the heart and is thus the major determinant of myocardial oxygen and energy consumption. By increasing wall thickness, ECM patch transplantation could relieve tension in the myocardial wall and subsequently reduce consumption of oxygen. An epicardial ECM patch may also modify the mechanical force vectors along the ventricular wall to favor an energetically more beneficial contraction. Furthermore, decreased wall tension might 1) increase perfusion to peri-infarct regions with barely surviving or stunned/hibernating cardiomyocytes and other cardiac cells through collateral vessels by relieving external pressure on thin-walled capillaries; and 2) produce mechanical signals for cardiomyocytes with prosurvival, and even proliferative, effects and 3) produce inhibitory signals to the active collagen deposition process, possibly sensed by activated myofibroblasts mechanistically. This process could explain the diminished fibrous scar content observed in our study in both patch groups. Others, including Mewhort et al. using a mouse model of MI (42), have also reported ECM patch-induced improved cardiac function. Further studies to assess the possible mechanisms underlying the profunctional effects of ECM patch transplantation are necessary.

Our results demonstrate that the therapeutic effects of the AAM patch are mediated synergistically by: 1) mechanical support with ventricular unloading by the ECM patch transplantation, as demonstrated by improved ECHO parameters, attenuated fibrosis and preservation of troponin T-positive myocardium in both acellular ECM as well as in AAM patch-treated groups; 2) enhanced cardiomyocyte contractility through expression of proteins associated with contractile apparatus and Ca^2+^ signaling in the AAM patch group; 3) enhanced reparative microenvironment through activated anaerobic glycolysis, angiogenesis and cell viability; 4) decreased oxidative stress and enhanced antioxidant protection by increased glutathione metabolism; 5) upregulated CM mitogenic matricellular POSTN; and 6) modulated inflammation. Moreover, in the myocardium of AAM patch-treated animals, the expression of TGM2 was greater than that in ECM patch-treated animals. This result suggests myocardial protection against ischemic injury as previously shown by Szondy et al. who demonstrated an increased myocardial injury after ischemia/reperfusion in TGM2^-/-^ knockout mice (43).

When evaluating the results of our study, it must be kept in mind that the samples were collected eight weeks after intervention opening the window for evaluation of true maintained therapeutic effect. This time-point, however, does not enable us to characterize the early events that take place during active inflammation or at the early phases of tissue destruction and fibrosis (44). Further research, especially targeting the earlier times of myocardial remodeling, is thus warranted to gain more in-depth understanding of the dynamics of healing induced by AAM patch. Nevertheless, it is plausible to state that the AAM patch transplantation has the ability to alter the pathological post-MI healing-by-scar dynamics by activating critical repair processes such as angiogenesis, cardioprotective and cardiogenic pathways, thus prolonging the reparative/restorative window or inducing a secondary healing response (27).

In conclusion, our results demonstrate that the epicardially transplanted AAMs-enclosing matrix patch significantly improves myocardial function after ischemic damage. The composite graft’s therapeutic effect is mediated by a combination of mechanical support, via ventricular unloading, and an activation of myocardium-protecting pathways, cardio-regenerative signaling, and angiogenesis. These results strengthen the therapeutic profile of epicardial AAM patch transplantation as a cost-effective, clinically feasible cellular adjuvant therapy for myocardial rescue and repair.

## Methods

A full-length description of the methods is available as an online-only data supplement (Detailed Materials and Methods) and Figure 1 illustrates the main parts of the study protocol and setting. Briefly, both atrial appendages were harvested and processed to AAMs from three male donor mice for each syngeneic transplant (42 mice in total). Forty age-matched male mice (homozygous 129X1/SvJ) were divided into the following groups: Sham-operated (Sham group, n=4); left anterior descending artery (LAD) ligation to induce MI (MI group, n=9); LAD ligation followed by epicardial transplantation of ECM patch without AAMs (ECM patch group, n=13); and LAD ligation followed by AAM patch, an ECM patch with AAMs, transplantation (AAM patch group, n=14). Cardiac function was evaluated weekly by echocardiography (ECHO), and after an 8-week follow-up, animals were sacrificed, and hearts collected for analyses.

Another set of mice was used to evaluate the mechanisms of AAMs. Male mice (strain 129X1/SvJ) were divided into three groups to receive either epicardial ECM (n=7) or AAM (n=7) patch therapy after MI or Sham control (n=4). As previously described, animals were sacrificed, and hearts were collected and cryopreserved for samples after the 8-week follow-up. Transplant, subtransplant and remote septal tissue samples were dissected from cryosections and were subsequently processed for label free site-targeted proteomic analysis (DIA-nanoLC-HD-MS^E^; AAM patch, n=4×3; ECM patch, n=4×3).

### Statistics

The statistical significance of column data presented in figures 2 to 5 was analysed using two-tailed Mann-Whitney U-test. Mann-Whitney U-test was chosen because the data was not normally distributed. The data normality was tested using Shapiro-Wilk test. Data derived from the ECHO follow up was analysed using ordinary one-way ANOVA.

### Study approval

The study conformed to the guide for the care and use of laboratory animals published by the US National Institutes of Health (NIH publication No. 85-23, revised 1996). Study protocols were approved by the Animal Experimentation Committee of the University of Helsinki, Finland, and the Provincial State Office of Southern Finland (ESAVI/8054/04.10.07/2016).

## Supporting information

Compiled Supplement File

Figure S2, Heatmap.

Datafile S1. All identified proteins and DEPs in locational order

## Author contributions

X.Y. and Mi. L. planned and performed experimental research, performed and supervised analyses, and wrote the article; J.T. planned and carried out immunohistochemistry, histomorphometry, image analysis, and wrote the article; V.S., Ma.L. and R.S. planned and performed proteomics analyses and bioinformatics, and wrote the article; E.M., M.K., A.H. and Z.Z. planned studies, provided critical resources and funding; E.K. planned and supervised the study, performed analyses, provided critical resources and funding, and wrote the article; all authors contributed to the writing and editing of the manuscript.

## Acknowledgments

The authors thank Päivi Leinikka for her invaluable help in carrying out the animal experiments, Lahja Eurajoki and Nada Bechara-Hirvonen for their expert laboratory assistance and Sole Lätti for the illustrations. Tissue processing was carried out at the Tissue Preparation and Histochemistry Unit of the Faculty of Medicine, University of Helsinki. Scanning of histological sections on slides was carried out at the Digital Microscopy and Molecular Pathology Unit, Institute for Molecular Medicine Finland (FIMM). Proteomics analyses were performed at the Meilahti Clinical Proteomics Core Facility, HiLIFE supported by Biocenter Finland.

## Disclosures

None for the authors. Antonio Graziano, a member of the AADC consortium, is the founder of and owns stock in HBW srl, manufacturer of the Rigenera HBW tissue processor.

**Correspondence and requests for materials** should be addressed to E.K.

---

**Figure S1.**
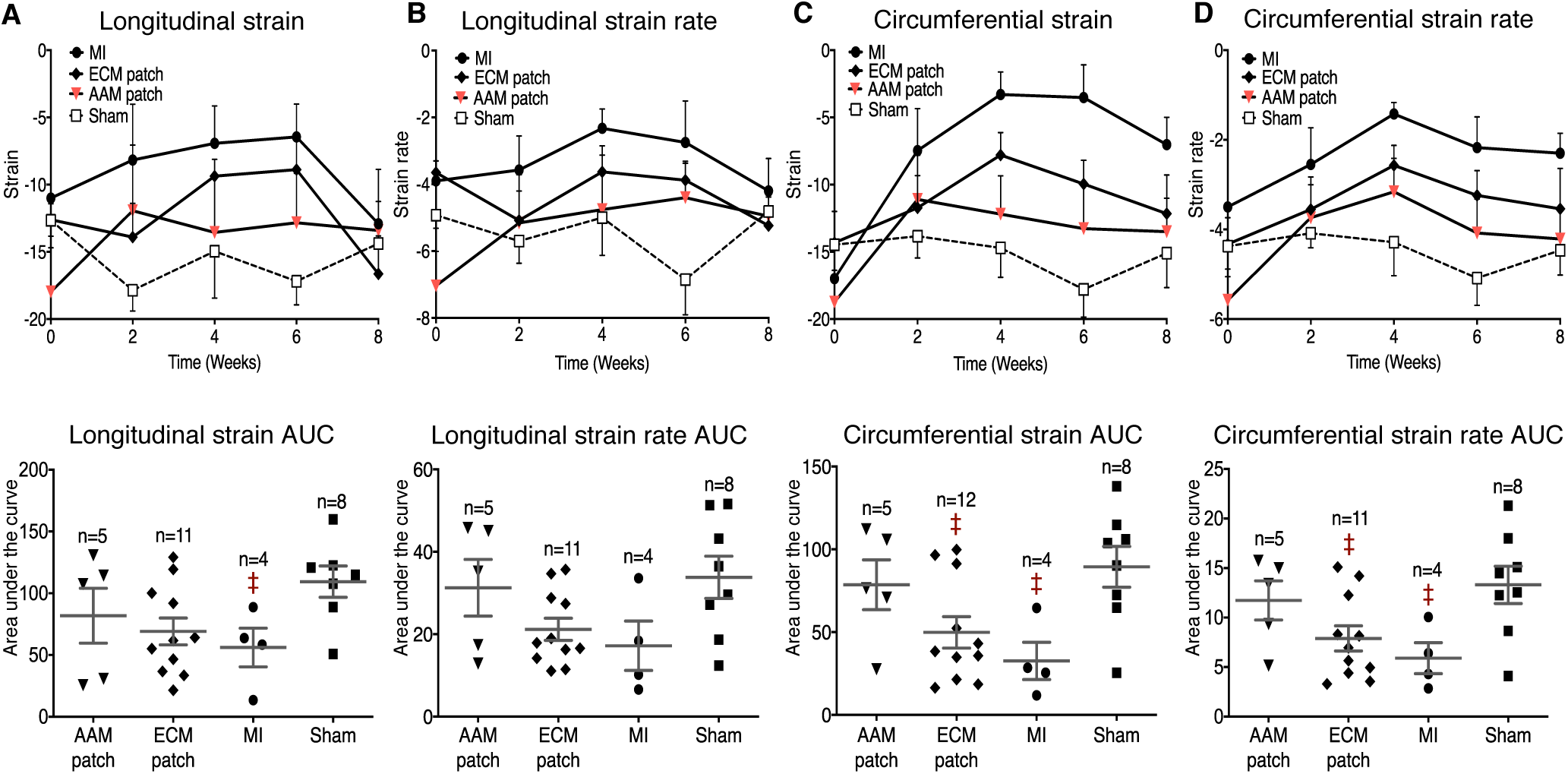
Complete data from the strain analysis. Data from the strain analysis performed during the 8-week follow-up period. The strain analysis was performed in 2-week intervals and was assessed in both the circumferential and longitudinal direction. The strain values are shown in upper half of the panels, and the corresponding AUC values below. Panels (**A**) and (**B**) show the values of the longitudinal strain and strain rate, respectively, while panels (**C**) and (**D**) present the circumferential strain and strain rate, respectively. Due to technical difficulties concerning the quality of the ECHO data, our analyses did not detect significant differences between groups in many cases. Data was assessed using unpaired Mann-Whitney *t*-test and normality of the data was asessed using Shapiro-Wilk test for normality. The scatter plot data represent mean ± SEM, while the n-values represent the number of mice measured. ‡*P* < 0.05; ‡, *P* vs. the Sham group.

