## Supplementary material for "Epicardial therapy with atrial appendage micrografts salvages myocardium after infarction": Compiled Supplement File

### Supplemental material

#### Detailed Materials and Methods

*Perioperative period*

To harvest atrial appendages, we anesthetized 8 to 9-week-old donor mice (homozygous males 129X1/SvJ, Laboratory Animal Center, University of Helsinki, Finland) with 4% isoflurane inhalation. Intubation was performed using a 20G needle tube. Maintenance of anesthesia was achieved with 2% isoflurane in a gas mixture consisting of 95% oxygen and 5% carbon dioxide. Ventilation was performed using a rodent ventilator (TOPO Dual Mode Ventilator, Kent Scientific Corp., Torrington, CT, USA) in pressure-control mode. Hearts were excised after thoracotomy, and a pool of both right and left atrial appendages were collected into cold cardioplegia to produce transplants in a similar manner as previously described (1).

Briefly, animals were sacrificed after tissue collection. To avoid material losses with the small sized atrial appendages, we used a Rigenera machine (Human Brain Wave Srl, Torino, Italy) to process the atrial appendages into micrografts (AAMs). Specifically, the atrial appendages were rinsed in ice-cold cardioplegia solution with removal of all extra adipose and connective tissue and cut into 1-3 $mm^3$ pieces. The pieces were placed on the Rigeneracon blade (Rigenera-system, HBW s.r.l., Turin, Italy) for mechanical isolation of AAMs. The blade is rotated with use of an electric drill at 100 rpm for 30s. The AAMs suspension is then collected with a syringe and the blade is rinsed with cardioplegia to collect all AAMs from the blade. The AAMs suspension was subsequently centrifuged at 400 g for 5 min and the supernatant was discarded. An extracellular matrix sheet (CorMatrix® ECM™ for cardiac tissue repair, CorMatrix Cardiovascular Inc., Roswell, GA, USA) was peeled into four layers, and 1-ply sheets were cut into circles using an 8 mm diameter tissue punch (Miltex™ Standard Biopsy Punches 762165, Integra LifeSciences Corp., Plainsboro, NJ, USA) and wetted in cardioplegia for 20 minutes. Then, the AAMs were divided into equal portions to obtain AAMs from approximately three right and three left appendages (from three donor mice) for each transplant. The AAMs were evenly dispersed onto sheet matrix and sealed with a thin layer

(10 µl volume) of Tisseel™ tissue glue (Baxter AG, Vienna, Austria). Premade AAM patches were kept on ice to prevent cell necrosis and to halt the cellular metabolism before transplantation. The operations were carried out in a blinded and randomized manner in relation to sheet type therapy with a separate person (Mi.L.) in charge of production, randomization and allocation of the patches to each animal. A separate person was in charge of the surgery and echocardiography (ECHO) (X.Y.). Our first set consisted of a total of 40 age-matched male mice (homozygous 129X1/SvJ) divided into four groups as follows: Sham-operated (Sham group, n=4), LAD ligation to induce MI (MI group, n=9), LAD ligation followed by epicardial ECM patch transplantation (ECM patch group, n=13), and LAD ligation followed by AAM+ECM patch transplantation (AAM patch group, n=14). Both atrial appendages were harvested and processed to AAMs from three male donor mice for each syngeneic transplant (42 donor mice in total). Animals were anesthetized using the method described above. Ligation of the coronary artery was performed from thoracotomy using an 8-0 Prolene™ suture (Ethicon, Johnson & Johnson Medical Devices, New Brunswick, NJ, USA). Overall mortality in the LAD ligation operation was 23.6% and the animals which survived the operation were randomised into the different interventional groups.

A premade transplant was fitted onto the recipient heart with the AAMs-containing side facing the epicardium of the left ventricle. In this model of LAD ligation, the site of visibly confirmed infarction and ischemia was the patch transplantation site. The AAM or ECM patch was further fixed against the epicardium by three sutures to ensure that the patches remained in place.

The recipient mice were followed for 8 weeks. Weekly ECHO measurements were performed using a Vevo 2100 Ultra High Frequency ultrasound system (Fujifilm VisualSonics Inc., Toronto, Canada). After the 8 weeks of follow-up, animals were sacrificed, and hearts were collected. The tissue samples obtained from the animals were taken from the ventricular level, apex, and atrium. Ventricular samples were fixed in paraformaldehyde solution (Sigma-Aldrich, St. Louis, MO, USA) for 48 hours and then preserved in 70% ethanol until embedded in paraffin. Paraffin-embedded sections were cut with a microtome and used in the Picrosirius Red assay and immunohistochemistry staining. Apex samples were immersed in RNAlater solution (AM7021, Thermo Fischer Scientific Inc., Waltham, MA, USA) and incubated overnight at +4°C before storage at -80°C for further use in a quantitative PCR assay to assess atrial natriuretic peptide (ANP) mRNA expression.

Another set of mice was treated to evaluate the AAM mechanism. Male mice (129X1/SvJ) were divided into two groups, namely, those receiving ECM patch therapy only and those receiving AAM patch therapy. A total of 21 donor appendages were used for generating the AAM patches (as detailed above) that were transplanted into 7 male mice. Induction of MI and transplantation of either ECM or AAM patches were performed as described above. After eight weeks of follow-up, animals were sacrificed, hearts were collected and ventricular tissue samples were snap frozen using liquid nitrogen and Tissue-Tek OCT® compound (Sakura Finetek Inc., Torrance, CA, USA). Cryosections of 12-µm thickness were cut from snap-frozen samples using a cryomicrotome (Leica CM3050S, Leica Biosystems Inc., Richmond, IL, USA). Cryosectioned samples were used for tissue proteomics analysis, which is explained more extensively later (See also Figure 1), and for validative qRT-PCR.

*Ethical aspects*

The study conformed to the guide for the care and use of laboratory animals published by the US National Institutes of Health (NIH publication No. 85-23, revised 1996). Study protocols were approved by the Animal Experimentation Committee of the University of Helsinki, Finland, and the Provincial State Office of Southern Finland (ESAVI/8054/04.10.07/2016).

*Evaluation of heart failure*

Functional evaluation of heart failure was performed over the 8-week follow-up period of the study by transthoracic ECHO using a visual sonic Vevo 2100 ultrasound device. Obtained ECHO data was analysed with Vevo LAB 1.7.1 software for conventional parameters, and supplemental strain analysis using plugin (Vevo Strain Analysis Software, Fujifilm VisualSonics Inc.). During the ECHO follow-up, left ventricular internal diameter (LVID), interventricular septal diameter (IVS), left ventricular posterior wall diameter (LVPW), and ventricular volume (LV Vol) were measured in the end systolic and end diastolic point. Fractional shortening (FS), left ventricular ejection fraction (LVEF), stroke volume (SV), heart rate (HR), and cardiac output (CO) were also included in the follow-up. LVID, IVS, LVPW, SV, HR, and CO were measured every 2 weeks, thus yielding five measurement points. EF, FS, and LV Vol were measured weekly from the start of the experiment, thus yielding nine measurement points. LVEF was measured as previously published (2). In addition to conventional ECHO follow-up parameters described above, myocardial strain analysis was performed to assess myocardial function over the follow-up period as a supplemental method. *Strain* assesses the myocardial wall movements, i.e., wall deformation. Strain can be measured by ECHO from the longitudinal and circumferential planes, and it offers additional information for evaluation of regional myocardial contractility with the capability to differentiate active and passive deformation, i.e., stretching and contraction. The negative strain values represent contraction, and *the strain rate* is the value of the strain as a function of time, i.e., how rapidly deformation occurs. Strain analyses were performed in 2-week intervals over the entire 8-week follow-up period. Circumferential strain, longitudinal strain, and corresponding strain rates were measured. In addition to LVEF, strain analysis is considered a relatively reliable method to quantify heart failure as well as conventional ECHO (3).

*Sample preparation for ANP mRNA expression*

RNA extraction was performed using tissue samples obtained from the apex region. Tissue RNA was extracted using 900 µl QIAzol Lysis Reagent (Qiagen NV, Venlo, The Netherlands) per 50 mg of tissue and a Precellys homogenizer (Bertin Technologies SAS, Montigny-le-Bretonneux, France) with CK28 Hard tissue homogenizing 2.8 mm ceramic beads (Bertin Technologies SAS). Samples were homogenized at 5500 rpm, 3 x 20 s with a 15 s break. After chloroform extraction, purification was continued with a Qiagen RNeasy extraction kit (Qiagen NV). Extracted RNA was then converted into complementary DNA (cDNA) with reverse transcriptase reaction. Quantitative PCR (qPCR) analysis using the Sybr Green method was then used to analyze the expression of the target ANP mRNA and the reference housekeeping gene 18S. The primers used were: ANP 5'-GAAAAGCAAACTGAGGGCTCTG-3' (forward) and 5'-CCTACCCCCGAAGCAGCT-3' (reverse) from Oligomer Oy (Helsinki, Finland). Primers for 18S were 5'-ACATCCAAGGAAGGCAGCAG-3' (forward) and 5'-TTTTCGTCACTACCTCCCCG-3' (reverse) from Sigma-Aldrich. Four 96-well PCR plates were used in this study for the analysis of ANP expression. The samples were analyzed in three different PCR plates; the fourth plate included calibrator samples from the three previous PCR plates to ensure that the values obtained from the different PCR plates were comparable to each other. The relative gene expression of each sample was calculated using the double delta Ct (ΔΔCt) method.

*Sample preparation for validative qRT-PCR (Postn, Myh7, Gpx3, Tgm2)*

RNA was extracted from cryosections of the myocardium from AAM patch (n=7), ECM patch (n=7) and Sham groups (n=4). For each sample, six 20µm-thick sections were collected and processed according to the RNA extraction kit manufacturer's instructions (NucleoSpin® RNA XS, Macherey-Nagel, Germany). Samples' RNA integrity number (RIN) values were between 8.0-9.5. Purified RNA was subsequently converted to cDNA and used for qRT-PCR. The primers for qRT-PCR were as follows: *Postn* 5'-GATAAAATACATCCAAATCAAGTTTGTTCG-3' (forward) and 5'-CGTGGATCACTTCTGTCACCGTTTCGC-3'(reverse) (4), *Myh7* 5'-ATCTTCTCCATCTCT-GAC-3'(forward) and 5'-ATTTGATCTTCCAGGGTA-3' (reverse), *Gpx3* 5'-CTGACAGAC-CAATACCTT-3' (forward) and 5'-GAATGACCAAGCCAAATG-3' (reverse), *Tgm2* 5'-AGGACATCAACCTGACCCTG-3' (forward) and 5'-CTTGATTTCGGGATTCTCCA-3' (reverse) (5) and primers for housekeeping 18S were identical as in ANP mRNA measurement. All primers were ordered from Sigma-Aldrich, primers for *Myh7* and *Gpx3* were designed using online design tool (OligoArchitect$^{tm}$ Online, Sigma-Aldrich). ΔΔCt method was used for calculating relative mRNA expression. Data was analyzed using unpaired and one-tailed parametric t-test with Welch's correction for non-identical SDs. Normality was tested using Shapiro-Wilk normality test.

*Tissue fibrosis*

Collagen-binding Picrosirius Red (ab150681, Abcam, Cambridge, UK) was used to evaluate the fibrosis and collagen content of the samples. Picrosirius Red staining was performed using a protocol previously described (7). By using a circularly polarized light, it is possible to quantify the amount of the different types of collagen fibers based on their thickness and birefringent appearance (6-8). Using Picrosirius Red staining and circularly polarized light, thinner type-III fibers appear green, whereas thicker and more mature type-I fibers appear red or yellow.

Seven images were obtained from the endocardial surface of the ventricle samples using 40x magnification and circularly polarized light using a Leica DMR light microscope, Leica DFC450 microscope camera and Leica PL APO 40x/0.85 corr object (Leica, Wetzlar, Germany). Imaging was performed during a single session using the same settings throughout. The images were obtained from the endocardial surface corresponding the center point of the infarction scar as this is considered the area most at risk during myocardial ischemia.

A total of 154 images were obtained from 22 animals. The collagen content of images was evaluated using Fiji image processing software (9) and macros generated by us. The amount of bright, orange, and green fibers was calculated, and total collagen content was measured by combining the amount of previously measured parameters. The tissue area in each picture was measured manually.

*Ventricular remodeling and histological tissue preservation*

Picrosirius Red-stained sections were scanned using normal bright-field light using an Epson Expression 1680 pro scanner (Epson, Nagano, Japan). The images obtained were used to measure the thickness of the ventricular wall and infarct transmurality. Each scanned slide included three sections, making a total of 78 scanned images from 26 animals (including four animals from the Sham group).

The thickness of the ventricular wall at the site of the infarction was manually measured at the following three points using Fiji image processing software: at the midpoint of the scar and halfway from the midpoint to infarction scar border on both sides. After the ventricular wall thickness at the infarction site was measured, the thickness of the collagenous part of the infarction scar was measured. Again, the measurements were performed manually using three different measurement points in the same fashion as described above.

The morphometry of the ventricular tissue samples was also assessed using planimetry. In both the AAM and ECM patch groups, we first measured the angle of epicardial fibrotic tissue, constituted by the transplanted patch with or without AAMs, reactive granulation tissue and fibrosis (transplant area). A second planimetric measurement from the AAM patch, ECM patch and MI groups included only the angle in which the fibrotic tissue extended from the epicardial surface to the deeper portions of the myocardium (scar area).

The tissue located inside the infarction scar and left unstained (nonfibrotic tissue) by Picrosirius Red staining was also compared between the AAM and ECM patch groups using the particle calculator function in Fiji image analysis software[9]. Using the FIJI particle calculator, we included nonfibrotic tissue nodules with an area greater than 100 pixels in our analysis. After the number of nonfibrotic tissue nodules was compared, we compared the total area of the nonfibrotic tissue nodules.

*Evaluation of the histological preservation of cardiac tissue*

The paraffin samples obtained from the ventricular level were stained using an anti-cardiac troponin T (cTnT) antibody (ab8295, Abcam plc., Cambridge, UK), Anti-Periostin antibody (ab14041, Abcam plc., Cambridge, UK), Anti-Glutathione Peroxidase 1 antibody (ab22604, Abcam plc., Cambridge, UK) and Anti-Transglutaminase 2 antibody (ab421, Abcam plc., Cambridge, UK).

The troponin T stained samples were scanned using a Pannoramic 250 Flash II slide scanner (3DHISTECH Ltd., Budapest, Hungary) with 40x and 20x bright-field objects. Scanned slides were then photographed using Pannoramic viewer software (3DHISTECH Ltd.). The area in the infarction scar was photographed using 40x magnification, and the area under the transplant (infarction scar, cell transplant, and vital myocardium under the infarction scar) was photographed using 20x magnification. In Sham group samples, 10 photographs were taken from the ventricular myocardium using 20x and 40x magnification. The Sham group images were used as a reference values when tissue troponin T content was analyzed. The background staining (for example, red blood cells) was manually reduced using Paint Shop Pro image photo editor software (Corel Corp., Ottawa, Canada). The immunohistological staining between the study groups in the infarction scar area and subtransplant area were analyzed with FIJI image analysis software (9, 10) using a macro generated by us. The background of the images was then also analyzed using a macro generated by us. The area of the troponin T-stained tissue was then adjusted taking the amount of the background area into account. Glutathione peroxidase 1 (GPX1), Periostin (POSTN) and Transglutaminase 2 (TGM2) stained sections were scanned using the same equipment as in the Troponin T staining described above. Representative and qualitative images were obtained and are presented in the article Figure 8A. Primary dilutions for POSTN, GPX1 and TGM2 were 1:500, 1:300 and 1:100, respectively.

*Sample preparation and quantitative label-free proteomics analyses*

A similar proteomics analysis workflow was utilized as reported by Lalowski et. al (11). Briefly, ventricular tissue samples of mouse hearts were snap frozen in Tissue-Tek OCT® compound. Next, 12-μm thick axial sections were cut from the ventricular regions with a cryostat (Leica CM3050S, Leica Biosystems Inc.). From each sample, 15 to 20 individual cryosections were collected depending on the overall size of the sample in an axial plane. The cryosections were fixed with ethanol, and Tissue-Tek OCT® compound was removed by rinsing with water.

One of the slides from each sample was stained with hematoxylin and eosin (H&E) to provide a more accurate visualization of the AAM or ECM patch in relation to the subtransplant tissue.

Based on direct visual assessment of the sections, supported with the visual aid of HE-stained sections from each sample, the sections were subcategorized into the following groups: 1) a transplant/patch group consisting of the AAM or ECM patch, reactive granulation tissue and fibrosis; 2) a subtransplant group consisting of the tissue predisposed to the ischemia right beneath the transplant to all the way to the adjacent luminal or endocardial border; and 3) a septal group consisting of the macroscopically vital looking myocardium from the endocardial side opposite to the subtransplant side all the way to the septal border of right ventricle. An overview of the process is presented in Figure 1.

Using clean sterile knives and a stereomicroscope (Leica EZ4HD, Leica Biosystems Inc.), the sections were cut in three parts as described above. The tissue was moved to corresponding Eppendorf tubes and stored on ice. Subsequently, 60 µl of lysis buffer (7 M Urea, 2 M Thiourea, 4% CHAPS) was introduced to each tube containing the tissue material. The mixture was then briefly centrifuged (E-Centrifuge, Wealtec Corp., Sparks, NV). Next, the tissue was vortexed and sonicated (Sonics Vibra-cell VC505, amplitude 25%, Sonics & Materials Inc., Newtown, CT) in pulses (one pulse lasting 1-2 s, 5 pulses total) with 5-minute intervals on ice after each pulse to ensure maximal solubilization of proteins. To complete the process of solubilization, the tubes were incubated on ice for 30 minutes. After centrifugation at 30,000 x g (Centrifuge 5417R, Eppendorf AG, Hamburg Germany), the supernatant was collected into two different tubes, with one for subsequent protein concentration measurement and one for data independent acquisition nanoscale high definition liquid chromatography coupled to tandem mass spectrometry (DIA-nanoLC-HD-MS$^{E}$) analysis and stored in -80°C.

The concentration of solubilized proteins was determined using industrial dye reagent concentrate (Cat#500-0006, Bio-Rad Laboratories Inc., Hercules, CA) and measured with a spectrophotometer (Victor™ X3 2030 multilabel reader, PerkinElmer LifeSciences Inc., Waltham, MA, USA).

We used 10 μg total protein for subsequent filter-assisted sample preparation (FASP) and data DIA-nano-LC-HD-MS$^E$ measurements as previously described (11, 12). Database searches were carried out against UniProtKB/Swiss-Prot reviewed mouse (release 2017_16956 entries) with the Ion Accounting algorithm and using the following parameters: peptide and fragment tolerance, automatic; maximum protein mass, 750 kDa; minimum fragment ions matches per protein, ⩾7; minimum fragment ions matches per peptide, ⩾3; minimum unique peptide matches per protein, ⩾2; primary digest reagent as trypsin; missed cleavages allowed, 2; fixed modification, carbamidomethylation C; variable modifications, deamidation (N, Q) and oxidation of methionine (M); and false discovery rate <4%.

The robust overall quality of the proteomic data was evaluated by drawing a heatmap from all of the quantified proteins (see Figure S2) confirming the site-specific clustering of the data according to the different study groups. Specifically, the heatmap was drawn with Heatmapper (www.heatmapper.ca) (13).

*Statistical methods*

For differential expression analysis (defining differentially expressed proteins, DEPs), the list was limited to those quantified with fold change, FC >1.5 and $P \leqslant 0.05$ by analysis of variance for all comparisons. The list of up/downregulated protein changes with their corresponding unique UniProtKB/Swiss-Prot identifiers served as inputs into Ingenuity® Pathway Analysis, IPA® bioinformatics analyses (Qiagen Bioinformatics, Redwood City, CA, USA). Unless otherwise stated, the results were analyzed using GraphPad Prism software (GraphPad Software Inc., La Jolla, CA, USA). Groupwise comparisons were evaluated using unpaired Mann-Whitney *t*-test, unless otherwise mentioned. *P*-values < 0.05 were considered statistically significant. Data are expressed as the mean±SEM.

**Members of AADC consortium (in alphabetical order) and their affiliations**

Antonio Graziano[1], Ari Harjula[2], Miia Holmström[3], Shengshou Hu[4], Tatu Juvonen[2], Esko Kankuri[5], Sari Kivistö[3], Markku Kupari[2], Mika Laine[2], Milla Lampinen[5], Jari Laurikka[6], Miia Lehtinen[2], Eero Mervaala[5], Tuomo Nieminen[7], Annu Nummi[2], Tommi Pätilä[8], Juha Sinisalo[2], Raili Suojaranta-Ylinen[9], Kari Teittinen[2], Antti Vento[2], Erika Wilkman[3], Yanbo Xie[4], Zhe Zheng[4]

1 Department of Public Health, Experimental Medicine and Forensics, University of Pavia, Italy.

2 Heart and Lung Center, Helsinki University Hospital, Finland.

3 HUS Medical Imaging Center, Radiology, University of Helsinki and Helsinki University Hospital, Finland.

4 National Clinical Research Centre of Cardiovascular Diseases, State Key Laboratory of Cardiovascular Disease, Fuwai Hospital, National Center for Cardiovascular Diseases, Chinese Academy of Medical Sciences and Peking Union Medical College, Beijing, People's Republic of China.

5 Department of Pharmacology, Faculty of Medicine, University of Helsinki, Finland.

6 Department of Cardiothoracic Surgery, Heart Center Co., Tampere University Hospital, Finland.

7 Helsinki University Hospital, University of Helsinki, Department of Internal Medicine, South Karelia Central Hospital.

8 Pediatric Cardiac surgery, Children`s Hospital, Helsinki University Hospital, Finland.

9 Department of Anesthesiology and Intensive Care, Helsinki University Hospital, Finland.

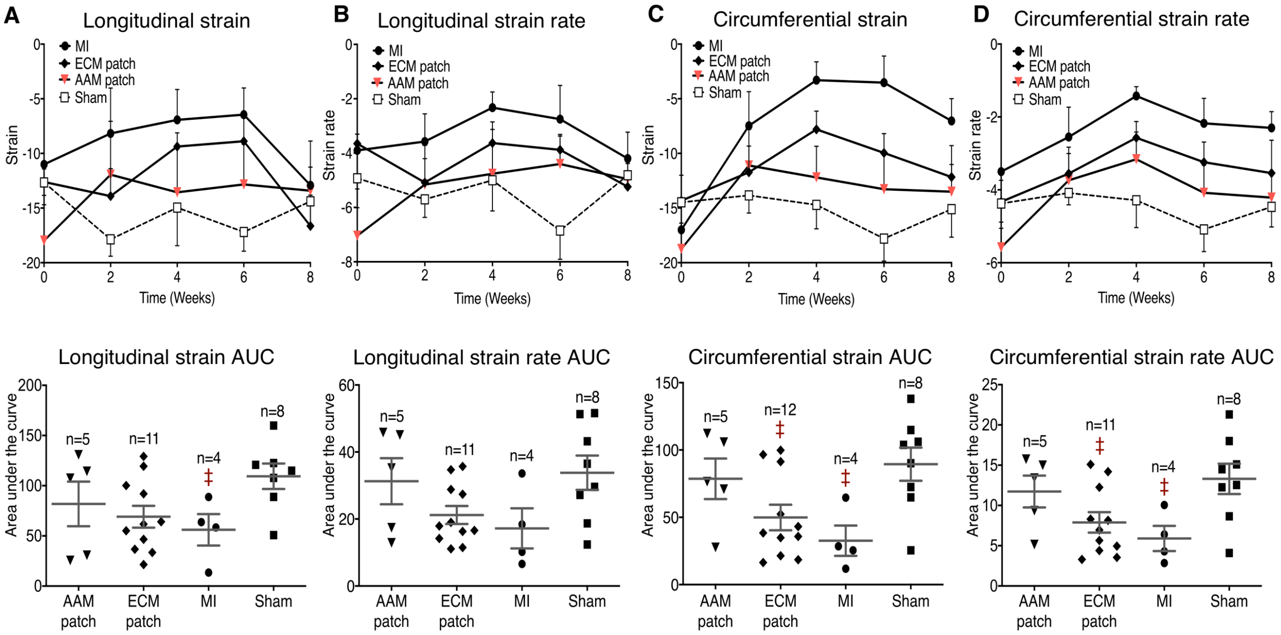

**Figure S1. Complete data from the strain analysis.**
Data from the strain analysis performed during the 8-week follow-up period. The strain analysis was performed in 2-week intervals and was assessed in both the circumferential and longitudinal direction. The strain values are shown in upper half of the panels, and the corresponding AUC values below. Panels (**A**) and (**B**) show the values of the longitudinal strain and strain rate, respectively, while panels (**C**) and (**D**) present the circumferential strain and strain rate, respectively. Due to technical difficulties concerning the quality of the ECHO data, our analyses did not detect significant differences between groups in many cases. Data was assessed using unpaired Mann-Whitney *t*-test and normality of the data was asessed using Shapiro-Wilk test for normality. The scatter plot data represent mean ± SEM, while the n-values represent the number of mice measured. ‡$P < 0.05$; ‡, $P$ vs. the Sham group.

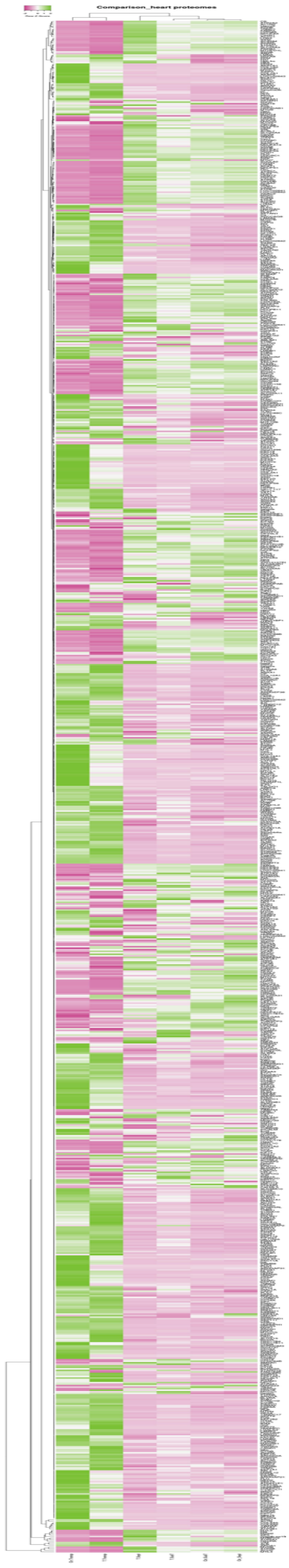

**Figure S2. Heatmap.**

Hierarchical clustering of proteomics data with protein and data grouping. An expression-based heat map was drawn utilizing all identified and quantified proteins (1005 unique IDs), average linkage and the Spearman rank correlation distance method. Ctr.Tranp- control transplant, T.Transp- treated transplant, Ctr.SubT- control subtransplant, T.SubT- treated subtransplant, Ctr.Sept- control septum, T.Sept- treated septum. The heat map was drawn with Heatmapper.

Due to the low resolution of the figure, we provide this Figure S2 as a separate file for more in-depth review alongside with the Data File S1 (Protemic Excel File).

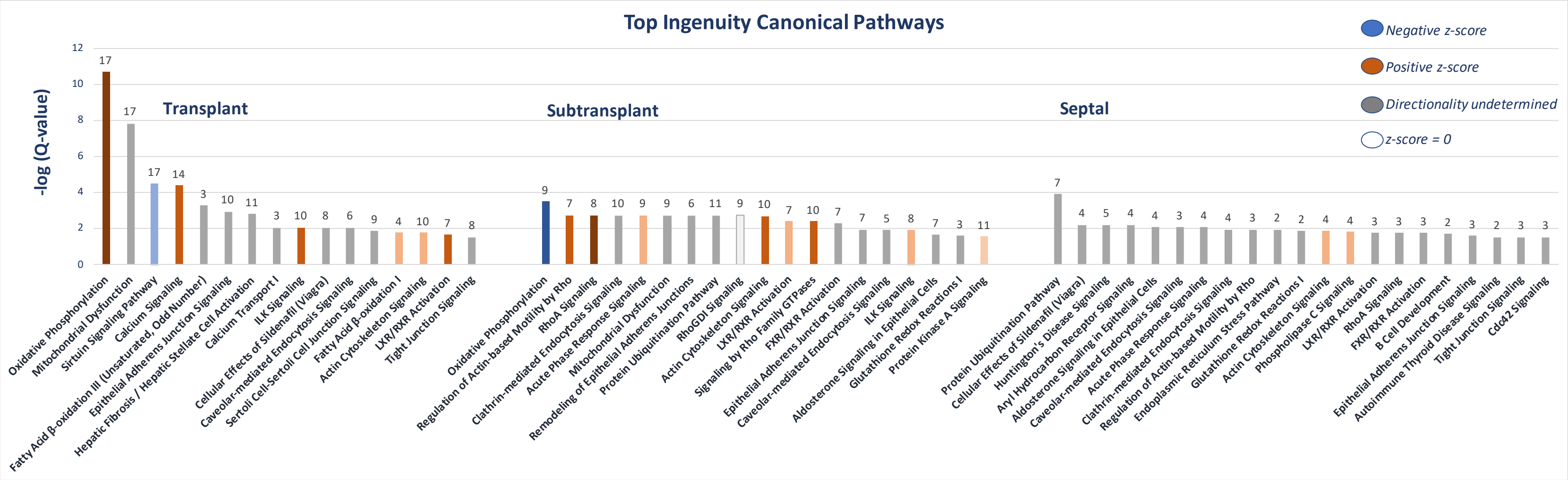

**Figure S3. All Top Ingenuity Canonical Pathways.**

Description of all the canonical pathways in each anatomical site as identified by IPA® software with given DEPs (lists of DEPs are provided in Online Data Supplement 2.4). The small counts at the top of each column represent the number of DEPs associated with a given pathway. The z-score indicates the direction of change in the pathway activity. The Q-value represents the B-H corrected *P*-value (Benjamini-Hochberg correction for *P*-value). -log (Q-value) ≥1.5 criterion for the level of significance was applied.

**Figure S4. All Diseases and Functions.**

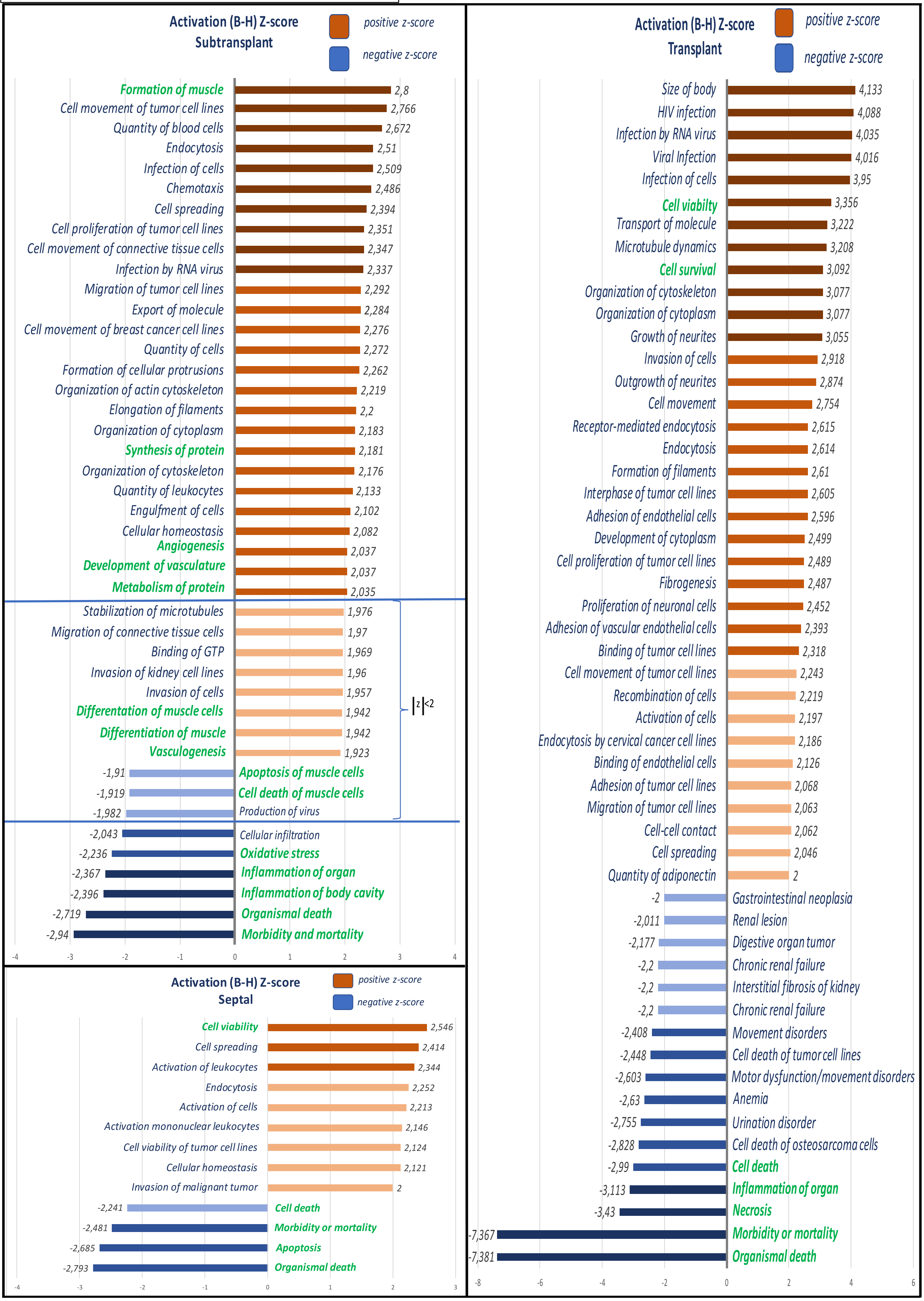

**Figure S4. All Diseases and Functions.**

Illustration of all the diseases and functions in each anatomical site identified by IPA® software with given DEPs (list of DEPs provided in Online Data Supplement 2.4). Functions labeled in green are shown in further detail in Figure 6**.** Small counts at the end of each column represents the z-score for each associated function or disease. |z-score| ≥ 2 was used as a criterion for statistical significance. B-H, Benjamini-Hochberg correction.

### Table S1. Comprehensive presentation of the ECHO data.

| | weeks | AAM patch | ECM patch | MI | Sham |
|---|---|---|---|---|---|
| **End diastolic left ventricular internal diameter (mm)** | 0 | 4.03±0.13 | 4.2±0.11 | 3.99±0.06 | 4.07±0.09 |
| | 2 | 4.9±0.29 | 4.71±0.18 | 4.9±0.34 | 4.23±0.05 |
| | 4 | 5.04±0.28 | 5.03±0.26 | 5.42±0.35 | 4.34±0.08 |
| | 6 | 5.02±0.28 | 4.73±0.22 | 5.45±0.36 | 4.2±0.13 |
| | 8 | 4.82±0.22 | 4.93±0.16 | 5.07±0.19 | 4.22±0.11 |
| **End systolic left ventricular internal diameter (mm)** | 0 | 2.92±0.16 | 3.09±0.15 | 2.76±0.09 | 3.27±0.12 |
| | 2 | 4.01±0.34 | 3.8±0.21 | 4.25±0.38 | 3.25±0.06 |
| | 4 | 4.14±0.34 | 4.34±0.31 | 4.96±0.39 | 3.61±0.1 |
| | 6 | 3.95±0.34 | 4.07±0.23 | 4.9±0.47 | 3.18±0.22 |
| | 8 | 3.86±0.33 | 4.07±0.2 | 4.39±0.3 | 3.12±0.12 |
| **End diastolic interventricular septal diameter (mm)** | 0 | 0.76±0.04 | 0.79±0.04 | 0.81±0.05 | 0.76±0.07 |
| | 2 | 0.85±0.06 | 0.77±0.06 | 0.74±0.09 | 0.71±0.03 |
| | 4 | 0.79±0.05 | 0.67±0.07 | 0.57±0.09 | 0.66±0.11 |
| | 6 | 0.85±0.04 | 0.88±0.06 | 0.74±0.12 | 0.75±0.06 |
| | 8 | 0.79±0.05 | 0.71±0.06 | 0.73±0.05 | 0.7±0.03 |
| **End systolic interventricular septal diameter (mm)** | 0 | 1.11±0.07 | 1.19±0.06 | 1.19±0.09 | 1.03±0.12 |
| | 2 | 1.2±0.1 | 1.06±0.08 | 1.02±0.14 | 1.02±0.1 |
| | 4 | 1.14±0.07 † | 0.9±0.11 | 0.7±0.1 | 0.93±0.13 |
| | 6 | 1.24±0.05 | 1.17±0.08 | 0.9±0.19 | 1.08±0.11 |
| | 8 | 1.13±0.09 | 0.98±0.09 | 0.97±0.09 | 0.99±0.06 |
| **End diastolic left ventricular posterior wall diameter (mm)** | 0 | 0.9±0.05 | 0.81±0.05 | 0.84±0.03 | 0.78±0.03 |
| | 2 | 0.7±0.06 | 0.8±0.06 | 0.86±0.09 | 0.8±0.02 |
| | 4 | 0.66±0.09 | 0.81±0.04 | 0.9±0.12 | 0.65±0.05 |
| | 6 | 0.82±0.05 | 0.83±0.06 | 0.77±0.1 | 0.86±0.06 |
| | 8 | 0.91±0.07 | 0.83±0.09 | 0.99±0.16 | 0.8±0.04 |
| **End systolic left ventricular posterior wall diameter (mm)** | 0 | 1.17±0.07 | 1.07±0.06 | 1.11±0.04 | 0.97±0.05 |
| | 2 | 0.87±0.09 | 1.05±0.08 | 1.02±0.12 | 0.97±0.02 |
| | 4 | 0.83±0.13 | 0.93±0.04 | 1.08±0.15 | 0.76±0.06 |
| | 6 | 1.05±0.06 | 0.94±0.07 | 0.93±0.13 | 1.08±0.07 |
| | 8 | 1.13±0.09 | 0.96±0.1 | 1.08±0.16 | 1.02±0.01 |
| **Left ventricular ejection fraction (%)** | 0 | 54.49±2.93 | 52.15±3.1 | 58.72±2.58 ‡ | 40.08±5.49 |
| | 1 | 36.69±2.56 † | 32.38±2.96 | 23.78±3.16 | 34.7±2.35 |
| | 2 | 39.27±4.31 | 39.84±3.65 | 32.12±3.82 | 46.5±1.71 |
| | 3 | 37.37±4.64 | 34.84±3.2 | 24.63±4.53 | 43.4±3.64 |
| | 4 | 38.73±4.25 † | 30.14±3.97 | 20.91±3.18 | 35.4±3.73 |
| | 5 | 42.07±5.25 † | 33.78±2.53 | 24.64±4.71 ‡‡ | 55.48±8.72 |
| | 6 | 45.18±3.9 † † | 33.43±2.9 | 23.3±6.49 ‡ | 49±4.81 |
| | 7 | 45.59±4.23 † † † | 38.13±2.47 † | 20.12±4.89 ‡ | 40.83±2.57 |
| | 8 | 43.91±4.91 | 36.68±3 | 28.92±6.09 | 51.73±2.29 |
| **Fractional shortening (%)** | 0 | 28.27±1.89 | 26.92±2.13 | 30.87±1.76 | 19.48±3.04 |
| | 1 | 17.65±1.34 † | 15.44±1.5 | 11.18±1.65 | 16.47±1.21 |
| | 2 | 19.53±2.38 | 19.74±2.08 | 15.4±1.95 | 23±1.01 |
| | 3 | 18.53±2.57 | 16.96±1.83 | 11.61±2.27 | 21.28±2.02 |
| | 4 | 19.21±2.31 † | 14.54±2.1 | 9.7±1.52 | 16.88±1.97 |
| | 5 | 21.65±3.35 | 16.28±1.37 | 11.71±2.37 ‡‡ | 29.58±6.02 |
| | 6 | 22.93±2.26 † † | 16.02±1.56 | 11.37±3.42 ‡ | 24.65±2.77 |
| | 7 | 23.21±2.6 † † † | 18.55±1.34 † | 9.46±2.41 | 19.8±1.47 |
| | 8 | 22.46±2.95 | 17.88±1.66 | 14.31±3.6 | 26.25±1.42 |
| **End diastolic ventricular volume (µL)** | 0 | 72.71±5.74 | 79.31±4.58 | 69.67±2.61 | 72.75±3.77 |
| | 1 | 90.93±7.89 † † | 97.77±8.88 † | 149.6±17.26 | 86.67±13.69 |
| | 2 | 119.8±16.49 | 105.2±9.84 | 123.7±22.37 | 79.75±1.89 |
| | 3 | 127.6±19.97 | 121.3±12.03 | 137.3±17.81 | 82.75±10.38 |
| | 4 | 127.1±17.54 | 125.5±16.83 | 158.2±22.44 | 85.25±3.73 |
| | 5 | 123.3±15.21 | 121.7±9.45 | 153.4±27.34 | 74.5±9.37 |
| | 6 | 123.5±18.57 | 107.9±12.51 | 151.2±22.72 | 79±5.79 |
| | 7 | 108.8±17.19 | 110.7±9.98 | 145.4±25.45 | 82.5±2.96 |
| | 8 | 111.8±11.61 | 116.3±9.01 | 124.1±9.84 | 80±5.12 |
| **End systolic ventricular volume (µL)** | 0 | 34.86±4.88 | 39.31±4.06 | 29.11±2.46 | 43.5±3.52 |
| | 1 | 59.71±7.29 † † | 68.92±9.43 † | 116.6±16.41 | 57±11.14 |
| | 2 | 79.69±15.85 | 65.54±8.79 | 88.89±21.16 | 42.5±1.76 |
| | 3 | 88.5±19.96 | 81.54±10.94 | 108.3±20.32 | 48±8.97 |
| | 4 | 84.58±17.49 | 92.62±16.61 | 130.1±23.3 | 55±3.67 |
| | 5 | 78.67±13.77 | 82±8.41 | 124±29.48 | 34.5±8.43 |
| | 6 | 74.17±17.35 | 74.33±10.88 | 124.6±25.4 | 41±7.22 |
| | 7 | 66.08±16.65 | 69.83±8.22 | 122.2±27.53 | 48.75±2.63 |
| | 8 | 68.67±12.31 | 75.75±8.47 | 92.11±11.46 | 38.75±3.75 |
| **Stroke volume (µL)** | 0 | 37.86±1.6 | 40±1.61 ‡ | 40.56±1.83 ‡ | 29.25±4.61 |
| | 2 | 40.01±3.28 | 39.63±3.32 | 34.87±3.76 | 37.03±1.59 |
| | 4 | 42.53±3.03 † | 32.98±3.46 | 28.11±2.49 | 30.3±3.76 |
| | 6 | 49.39±2.88 ** † † † | 33.58±2.88 | 26.67±4.86 | 37.75±1.72 |
| | 8 | 43.17±1.62 † | 40.58±2.42 | 32±3.64 | 41.25±2.78 |
| **Heart rate (BPM)** | 0 | 349±10.25 | 333.9±8.29 | 324.9±14.14 | 368.3±17.2 |
| | 2 | 373.7±7.48 | 372.7±16.62 | 370.1±12.49 | 324.8±23.13 |
| | 4 | 352.7±13.04 | 329.5±9.89 | 346±13.12 | 341.3±15.66 |
| | 6 | 356.6±9.04 | 368.9±12.08 | 373.3±5.56 | 371.5±17.33 |
| | 8 | 357.4±9 | 363.3±11.37 | 377.6±17.41 | 367.5±19.6 |
| **Cardiac output (mL/min)** | 0 | 13.21±0.74 | 13.31±0.67 | 13.11±0.79 | 10.75±1.32 |
| | 2 | 14.96±1.27 | 14.42±0.97 | 13.05±1.69 | 12±0.9 |
| | 4 | 14.88±1.05 * † | 10.78±1.1 | 9.95±1.13 | 10.18±0.89 |
| | 6 | 17.78±1.34 * † † | 12.23±0.86 | 9.98±1.83 | 14.09±1.17 |
| | 8 | 15.38±0.64 | 14.82±1.07 | 11.89±1.25 | 15.28±1.8 |

**Table S1. Comprehensive presentation of the ECHO data.** Comprehensive mean ECHO follow-up values at each time point of the 8-week follow-up. */†/‡*P* < 0.05; **/††/‡‡*P* < 0.002; ***/†††/‡‡‡*P* < 0.0002; *, *P* when compared against ECM patch group; †, *P* when compared against the MI group; ‡, *P* when compared against the Sham group. Groupwise analysis was carried out using one-way ANOVA.

| Table II, all associated DEPs for all canonical pathways and diseases and functions presented in Figure 6 | | |
|---|---|---|
| **Canonical pathways** | | |
| **Location** | **Pathway** | **Associated DEPs** |
| TRANSPLANT | | |
| | Oxidative Phosphorylation | Cox6c, NDUFS7, Cyct, NDUFB5,ATP5PO, MT-ND4, MT-CO2, NDUFV2, UQCR10, NDUFA6, CYC1, COX5A, NDUFS6, ATP5MF, NDUFS2, NDUFA3, UQCRQ |
| | Sirtuin Signaling Pathway | SLC25A4, NDUFS7, TRIM28, NDUFB5, TUBA4A, MT-ND4, PGAM2, ACLY, NDUFV2, TUBA8, NDUFA6, CYC1, NDUFS6, NDUFS2, NDUFA3, GOT2, SLC25A5 |
| | Calcium Signaling | MYH10, TNNC2, ATP2B1, ATP2A3, TPM3, MYH7, MYH11, Tpm2, MYL1, ATP2A1, CAMK2D, HDAC7, ACTA1, MYH1 |
| | Fatty Acid β-oxidation I | Eci3, ECI2, HADHA, ECI1 |
| | ILK Signaling | MYH10, FLNC, SH2B2, ILK, VIM, MYH7, MYH11, ACTA1, MYL1, MYH1 |
| | Actin Cytoskeleton Signaling | KNG1, MYH10, IQGAP2, CYFIP1, MYH7, MYH11, ACTA1, MYL1, MYL12A, MYH1 |
| SUBTRANSPLANT | | |
| | Oxidative Phosphorylation | ATP5PF, MT-ND5, NDUFB11, Cyct, ATP5MF, NDUFS6, MT-ND4, MT-CO2, UQCRQ |
| | Regulation of Actin-based Motility by Rho | PFN1, ARPC1B, MYL6, ACTB, ARHGDIA, GSN, MYL1 |
| | RhoA Signaling | PFN1, ARPC1B, SEPT9, MYL6, ACTB, SEPT11, GNA13, MYL1 |
| | Actin Cytoskeleton Signaling | PFN1, FN1, ARPC1B, MYL6, ACTB, MYH7, ACTN4, GNA13, GSN, MYL1 |
| | ILK Signaling | FN1, MYL6, ACTB, SH2B2, VIM, MYH7, ACTN4, MYL1 |
| | Glutathione Redox Reactions I | GPX3, GSTM2, GPX1 |
| SEPTAL | | |
| | Glutathione Redox Reactions I | GPX3, GSTM2 |
| | Actin Cytoskeleton Signaling | MYL6, MYH7, ACTA1, MYL12A |
| **Diseases and functions** | | |
| **Location** | **Disease or function** | **Associated DEPs** |
| TRANSPLANT | | |
| | Cell viability | ACLY, ADIPOQ, APOB, B2M, CAMK2D, CD47, COL4A3, DAB2, DPP3, EEF2, EIF3A, EIF3E, EIF4A1, ENO1, EPS15L1,FLI1, Gm21596/Hmgb1, HDGF, HLAA, HSPB1, HYOU1, ILK, ITGAM, KIF5A, KRT19, MYH11, MYO18A, NEFH, NT5C3A, PRPF19, PRPF8, PRPH, PSAP, PSMA4, RAB5A, RACK1, RAD50, SH3KBP1, SNW1, STAT1, TF, TRIM28, UNC119, VTN, XDH |
| | Cell survival | ACLY, ADIPOQ, APOB, B2M, CAMK2D, CD47, COL4A3, DAB2, DPP3, EEF2, EIF3A, EIF3E, EIF4A1, ENO1, EPS15L1, FLI1, Gm21596/Hmgb1, HDGF, HLA-A, HSPA4, HSPB1, HYOU1, ILK, ITGAM, JUP, KIF5A, KRT19, MYH11, MYO18A, NEFH, NT5C3A, PRPF19, PRPF8, PRPH, PSAP, PSMA4, RAB5A, RACK1, RAD50, SH3KBP1, SNW1, STAT1, TF, TRIM28, UNC119, VIM, VTN, XDH |
| | Cell death | ACLY, ADD1, ADIPOQ, ALDH3B1, ALOX15, AP2M1, APOB, ARL6IP5, ATP1A1, ATP2A1, ATP2B1, ATP4A, B2M, C4A/C4B, CA3, CAMK2D, CCNE1, CCT2, CCT3, CCT6A, CCT7, CCT8, CD36, CD47, CENPE, CHGB, COL4A2, COL4A3, COQ8A, COQ9, COX5A, Cyct, DAB2, DDX4, DLST, EEF2, EIF3B, EIF3E, EIF4B, ENO1, EPHX1, Ewsr1, FLI1, FXR1, Gm21596/Hmgb1, GNAS, GPLD1, HADHA, HDGF, HLA-A, HSPA4, HSPB1, HSPH1, HYOU1, ILF3, ILK, IQGAP2,ITGAM, JUP, KIF5A, KNG1, KRT19, LAMA2, LGMN, MARC2, MT-ND4, MTCH2, Mug1/Mug2, MYH10, MYH11, MYO18A, NASP, NCL, NDUFV2, NEFH, NT5C3A, NUMA1, PABPC1, PDLIM7, PRPF19, PRPF8, PRPH, PSAP, PSMA4, PSMB8, PYGL, Pzp, RAB1A, RAB5A, RACK1, RAD50, Rpl23a, RPL5, RPL7, SGCA, SH3KBP1, SLC25A4, SLC25A5, SNW1, SPTA1, SPTB, STAT1, TF, TFAM, TGFBI, TMED10, TNRC6A, TPM3, TRIM28, UNC119, VIM, VPS35, VTN, XDH, YWHAE, YWHAG, YWHAQ |
| | Inflammation of organ | ACADVL, ADIPOQ, ALOX15, APOA4, APOB, ATP1A1, ATP4A, B2M, C4A/C4B, CA3, CD36, CD47, COL3A1, COL4A3, EIF3E, ENO1, FXR1, Gm21596/Hmgb1, GSTK1, HLA-A, ITGAM, JUP, KNG1, KRT1, KRT16, LGMN, Mug1/Mug2, MYH10, PSAP, Pzp, RACK1, SCN1B, SGCA, STAT1, TF, TPM3, TUBA4A, TUBA8, VIM, XDH |
| | Necrosis | ACLY, ADIPOQ, ALDH3B1, ALOX15, AP2M1, APOB, ATP1A1, ATP2A1, ATP2B1, ATP4A, B2M, CA3, CAMK2D, CCNE1, CCT2, CCT3, CCT6A, CCT7, CCT8, CD36, CD47, CENPE, CHGB, COL4A2, COL4A3, COQ9, COX5A, Cyct, DAB2, DLST, EEF2, EIF3B, EIF3E, EIF4B, ENO1, EPHX1, FLI1, Gm21596/Hmgb1, GNAS, HADHA, HDGF, HLA-A, HSPA4, HSPB1, HSPH1, HYOU1, ILK, IQGAP2, ITGAM, JUP, KNG1, LAMA2, LGMN, MARC2, MT-ND4, MTCH2, Mug1/Mug2, MYH10, MYH11, NASP, NCL, NDUFV2, NEFH, PABPC1, PDLIM7, PRPF19, PRPF8, PRPH, PSAP, PSMA4, PSMB8, PYGL, Pzp, RACK1, RAD50, Rpl23a, RPL5, RPL7, SGCA, SH3KBP1, SLC25A4, SLC25A5, SNW1, STAT1, TF, TFAM, TGFBI, TMED10, TRIM28, UNC119, VIM, VTN, XDH, YWHAE, YWHAG, YWHAQ |
| | Morbidity or mortality | ACADVL, ACLY, ACSS1, ACTA1, ADD1, ADIPOQ, AFF4, ALOX15, AP2M1, APOB, ATP12A, ATP1A1, ATP2A1, ATP2B1, ATP5MF, B2M, C4A/C4B, CAPZA2, CCNE1, CD36, CD47, CENPE, COL3A1, COL4A2, COL4A3, COQ9, CSRP1, DAB2, EIF4A1, FLI1, FLNC, FXR1, Gm21596/Hmgb1, GNAS, GSTK1, HADHA, IQGAP2, ITGAM, JUP, KIF5A, KIF5B, KRT1, KRT19, LAMA2, Mug1/Mug2, MYH10, MYH11, MYH7,NASP, NUMA1, OAT, PRPF19, PRPF8, PSAP, Pzp, RAB5A, RAB6A, RAD50, SCN1B, SNAP23, SNX1, SRGAP3, STAT1, TF, TFAM, TGFBI, TMED10, TMOD1, TPM3, TRIM28, VIM, XDH, YWHAE, HDAC7, HLA-A, HYOU1, ILF3, ILK |
| | Organismal death | ACADVL, ACLY, ACSS1, ACTA1, ADD1, ADIPOQ, AFF4, ALOX15, AP2M1, APOB, ATP12A, ATP1A1, ATP2A1, ATP2B1, ATP5MF, B2M, C4A/C4B, CAPZA2, CCNE1, CD36, CD47, CENPE, COL3A1, COL4A2, COL4A3, COQ9, CSRP1, DAB2, EIF4A1, FLI1, FLNC, FXR1, Gm21596/Hmgb1, GNAS, GSTK1, HADHA, HDAC7, HYOU1, ILF3, ILK, IQGAP2, ITGAM, JUP, KIF5A, KIF5B, KRT1, KRT19, LAMA2, Mug1/Mug2, MYH10, MYH11, MYH7, NASP, NUMA1, OAT, PRPF19, PRPF8, PSAP, Pzp, RAB5A, RAB6A, RAD50, SCN1B, SNAP23, SNX1, SRGAP3, STAT1, TF, TFAM, TGFBI, TMED10, TMOD1, TPM3, TRIM28, VIM, XDH, YWHAE |
| SUBTRANSPLANT | | |
| | Cellular infiltration | ACTB, ALB, CCDC88A, CRYAB, ENO1, FABP4, FCGR2B, FN1, Gm21596/Hmgb1, HLA-A, Cellular infiltration HNRNPAB, HSPB1, LGALS1, PTPN6, SERPINA1, TGM2 |
| | Oxidative stress | GPX1, Gsta4, HSPB1, QDPR, SERPIND1 |
| | Inflammation of organ | ACTB, ACTN4, AHCY, ALB, APOA2, APOB, ARHGDIA, ATP4A, B2M, CKM, CORO1A, CROCC, DDX5, DPYSL2, ENO1, FABP4, FCGR2B, FN1, Gm21596/Hmgb1, GPX1, GSN, HLA-A, HNRNPAB, HPX, KRT15, KRT17, LAMB2, LGALS1, Mug1/Mug2, POSTN, PSAP, PTPN6, SERPINB1, TF, TGM2, VIM |
| | Inflammation of body cavity | ACTB, ALB, APOA2, APOB, ATP4A, B2M, CKM, CROCC, DDX5, ENO1, FCGR2B, FN1, Gm21596/Hmgb1, GPX1, HLA-A, HNRNPAB, HPX, LGALS1, Mug1/Mug2, PTPN6, SERPINB1, TF |
| | Organismal death | ACTB, ACTN4, ALDH1A1, APOB, ARHGDIA, ARHGDIB, ATP2B1, ATP5MF, B2M, CLIC4, CPT1B, CRYAB, CTTN, CYGB, DDX5, DES, DNM2, ERC1, FCGR2B, FN1, Gm21596/Hmgb1, GNA13, GNAS, GPX1, GPX3, GSN, HADHA, HNRNPAB, HNRNPD, HSPB6, HSPE1, IDH3A, LAMB2, LAMP2, LCP1, LMNB2, LRP1, LUM, MAP, Mug1/Mug2, MYH7, MYL6, PFN1, POSTN, PRPF19, PRPF8, PSAP, PTPN6, QDPR, S100A13, SEPT9, SERPINA1, SERPINA3, SERPIND1, SERPINH1, SLC9A3R1, SNX1, SRGAP3, TF, TGM2, TMOD3, VIM |
| | Morbidity or mortality | ACTB, ACTN4, ALDH1A1, APOB, ARHGDIA, ARHGDIB, ATP2B1, ATP5MF, B2M, CLIC4, CPT1B, CRYAB, CTTN, CYGB, DDX5, DES, DNM2, ERC1, FCGR2B ,FN1,Gm21596/Hmgb1, GNA13, GNAS, GPX1, GPX3, GSN, HADHA, HLA-A, HNRNPAB, HNRNPD, HSPB6, HSPE1, IDH3A, LAMB2, LAMP2, LCP1, LMNB2, MYH7, MYL6, PFN1, POSTN, PRPF19, PRPF8, PSAP, PTPN6, QDPR, S100A13, SEPT9, SERPINA1, SERPINA3, SERPINB1, SERPIND1, SERPINH1,SLC9A3R1, SNX1, SRGAP3, TF, TGM2, TMOD3, VIM, LRP1, LUM, MAP4, Mug1/Mug2 |
| | Apoptosis of muscle cells | CAMK2D, CRYAB, EEF1D, GNAS, GPX1, GSN, HSPB1, HSPB6, HSPE1, MTPN, S100A6 |
| | Cell death of muscle cells | CAMK2D, CRYAB, EEF1D, GNAS ,GPX1, GSN, HADHA, HSPB1, HSPB6, HSPE1, MTPN, NCL, S100A6 |
| | Differentiation of muscle | ALB, DDX5, DES, EIF5A, FN1, GPX1, LGALS1, MTPN, MYH7, OGN, SNW1, SPAG9, TGM2, TRIM72 |
| | Differentiation of muscle cells | ALB, DDX5, DES, EIF5A, GPX1, LGALS1, MTPN, MYH7, OGN, SNW1, SPAG9, TGM2, TRIM72 |
| | Vasculogenesis | APOB, ARHGDIA, CLIC4, CRYAB, CYGB, DNM2, FABP4, FN1, Gm21596/Hmgb1, GNA13, GPX1, GSN, HECTD1, IDH3A, LGALS1, LRP1, PSAP, PTPN6, SEPT9, SERPINA3, SERPIND1, SERPINH1, TF, TGM2, VIM, YARS, YWHAZ |
| | Development of vasculature | ACTN4, APOB, ARHGDIA, CLIC4, CRYAB, CTTN, CYGB, DNM2, FABP4, FN1, Gm21596/Hmgb1, GNA13, GPX1, GSN, HECTD1, HSPB1, HSPB6, IDH3A,LGALS1, LRP1, LUM, Mug1/Mug2, NCL, NID1, PSAP, PTPN6, SEPT9, SERPINA3, SERPIND1, SERPINH1, TF, TGM2, VIM, YARS, YWHAZ |
| | Angiogenesis | APOB, ARHGDIA, CLIC4, CRYAB, CYGB, DNM2, FABP4, FN1, Gm21596/Hmgb1, GNA13, GPX1, GSN, HECTD1, HSPB1, HSPB6, IDH3A, LGALS1, LRP1, NCL, PSAP,PTPN6, SEPT9, SERPINA3, SERPIND1, SERPINH1, TF, TGM2, VIM, YARS, YWHAZ |
| | Metabolism of protein | APOA2, APOB, CCT4, CCT8, CRYAB, DDX39B, EIF4G3, EIF5A, FCGR2B, FN1, GPLD1, GSN, HNRNPD, HSPA1A/HSPA1B, HSPB1, Ighg2a, KRT17, LAMP2, LRP1, MTPN, NCL, PFN1, PSMD11, RNPEP, RPL5, SERPINA1, SERPINB1, SNX1, SNX3, TF, TGM2, VIM |
| | Synthesis of protein | DDX39B, EIF4G3, EIF5A, FCGR2B, FN1, GSN, HNRNPD, HSPA1A/HSPA1B, HSPB1, KRT17, MTPN, NCL, RPL5, VIM |
| | Formation of muscle | ACTN4, CAMK2D, CRYAB, DES, DNM2, FN1, GSN, KRT17, LAMB2, MYH7, MYL6, TGM2, TRIM72, VIM |
| SEPTAL | | |
| | Cell death | ALB, ANXA5, APOA1, CAPN2, CRYAB, CYB5A, EHD4, HLA-A, HSP90B1, HSPA5, HSPB1, Ighg2a, LGALS1, LRP1, LUM, PHB2, POSTN, PPIA, SERPINA3, SNX1, TAGLN2, TF, TGM2, TMOD3, UBE2N, YWHAB |
| | Morbidity or mortality | ACTA1, APOA1, CAPN2, CRYAB, EHD4, GPX3, HLA-A, HSP90B1, HSPA5, LRP1, LUM, MYH7, MYL6, PHB2, POSTN, PPIA, SERPINA3, SLC9A3R1, SNX1, TF, TGM2, TMOD3, UBE2N |
| | Apoptosis | ALB, ANXA5, APOA1, CAPN2, CRYAB, CYB5A, EHD4, HSP90B1, HSPA5, HSPB1, LGALS1, LRP1, LUM, PHB2, PPIA, SERPINA3, SNX1, TAGLN2, TF, TGM2, TMOD3, YWHAB |
| | Organismal death | ACTA1, APOA1, CAPN2, CRYAB, EHD4, GPX3, HSP90B1, HSPA5, LRP1, LUM, MYH7, MYL6, PHB2, POSTN, PPIA, SERPINA3, SLC9A3R1, SNX1, TF, TGM2, TMOD3, UBE2N |
| | Cell viability | ALB, ANXA5, CAPN2, EHD4, HLA-A, HSP90B1, HSPA5, HSPB1, LRP1, PHB2, POSTN, PPIA, SERPINA3, TF, TGM2 |

**Table S2. All associated DEPs for all diseases and functions presented in Figure 6.**

List of all canonical pathways and diseases and functions presented in Figure 6 with all associated DEPs. As the number of associated DEPs for certain functions is vast, the same background color was used to group adjacent rows that belong to the same function.

**Datafile S1. All identified proteins and DEPs in locational order.**

The tables in question are available from the authors as a separate excel file. All identified proteins (UPs $\geq 2$) and corresponding DEPs (FC > 1.5, $p < 0.05$, UPs $\geq 2$) for each location are presented in the consecutive sheets, labeled as follows: List_all_*, all identified proteins; IPA_*, all DEPs; Transp, Transplant area; Stransp, subtransplant area; Sept, septal area; AAMs, AAMs patch group and ECM, ECM patch group. FC, fold change; p, p-value by ANOVA; UPs, unique peptides.

The proteomic data have been uploaded to the Mass Spectrometry Interactive Virtual Environment (MassIVE) for in-depth review (MSV000084120). The login information for accessing the data will become public when the manuscript is accepted for peer-reviewed journal. However, the corresponding author can be approached for specific questions concerning the raw data before the final publication.
