## Supplementary figures and images for "Epicardial therapy with atrial appendage micrografts salvages myocardium after infarction"

### Figure S2, Heatmap.

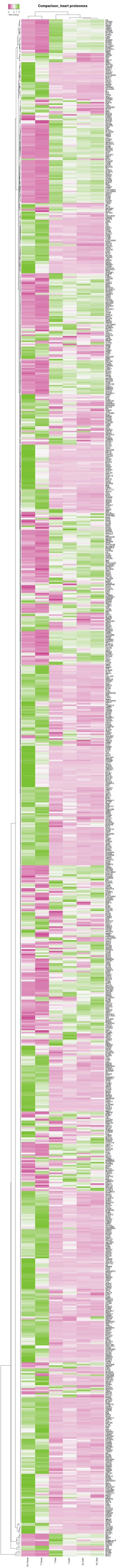
